# Copper transport to mitochondria by SLC25A3 contributes to skeletal myoblast differentiation and is required for survival of differentiated myotubes

**DOI:** 10.64898/2026.06.03.729837

**Authors:** AM Perez, JD Fivush, B Cordill, N Ferguson, Y Zhang, AT Mezzell, U Mattam, O Chaudhry, K Porter, S Maadaadi, D Secic, ME Bischoff, K Chella Krishnan, RA Kovall, JT Cunningham, MF Czyzyk-Krzeska, KE Vest

**Author notes:** Corresponding author: Katherine E. Vest.

## Abstract

Differentiation of skeletal muscle is associated with increased mitochondrial biogenesis and reliance of oxidative phosphorylation (OXPHOS). The terminal enzyme complex in the electron transport chain, cytochrome *c* oxidase (COX), requires copper for its assembly and activity, and copper delivery to mitochondria is essential for OXPHOS. However, when mitochondrial copper becomes essential during skeletal myoblast differentiation is not known. Here, we show that genetic deficiency of the mitochondrial copper and phosphate carrier SLC25A3 induced prior to myoblast differentiation leads to the formation of smaller myotubes, but SLC25A3 deficiency induced in mature myotubes leads to cell death and detachment. Both phenotypes are recapitulated upon genetic knockdown of COX17, a critical assembly protein for both COX copper cofactors, or by chemical inhibition of COX. Importantly, myotube death caused by SLC25A3 deficiency is rescued by copper supplementation or expression of an SLC25A3 variant that transports copper but not phosphate. Taken together these data support a model wherein copper transport by SLC25A3 and copper delivery to COX is critical for survival in mature myotubes.

## Introduction

Copper is a trace nutrient that contributes to fundamental cellular pathways like energy production, extracellular matrix modification, and pro-proliferative signaling in mammalian cells (1, 2). Despite its essential functions, excess or misallocated copper can be toxic due to redox imbalance, mis-metalation of other metalloproteins, or cuproptosis, a programmed cell death pathway initiated by excess copper in mitochondria (2). Despite the potential for mitochondrial copper to initiate cell death, copper delivery to mitochondria is essential due to its function as a cofactor in cytochrome *c* oxidase (COX), complex IV in the electron transport chain (ETC) that is required for ATP production by oxidative phosphorylation (OXPHOS) (3). The COX complex consists of 14 nuclear and mitochondrial-encoded subunits and requires more than 30 nuclear-encoded accessory factors for assembly (4). Of these assembly factors, several in the intermembrane space (ISM) are responsible for loading the copper A (Cu_A_) and B (Cu_B_) cofactors into the catalytic COX2 and COX1 subunits, respectively (5). The IMS copper chaperones include COX17, SCO1, SCO2, COA6, COX11, and COX19 (6–11). Individuals expressing pathogenic variants in these and other COX assembly factors exhibit a collection of symptoms including lactic acidosis, encephalopathy, and skeletal and cardiac muscle impairment (12). The copper used by COX is stored in the mitochondrial matrix as a labile pool and must be transported back to the IMS for incorporation into COX (13, 14). Thus, copper transport into the mitochondrial matrix is critical for the function of the ETC.

Copper is delivered into the mitochondrial matrix by SLC25A3, a member of the mitochondrial carrier family (MCF) proteins, a highly conserved group that transport metabolites and ions primarily into and out of the mitochondrial matrix (15–17). Although originally identified as a phosphate carrier, loss of SLC25A3 in multiple mammalian cell lines leads to decreased COX activity that can be rescued by copper supplementation (18–20, 15, 16). These genetic studies, coupled with in vitro transport assays, suggest that SLC25A3 also transports copper and contributes to COX assembly and activity. The *SLC25A3* gene encodes two isoforms alternatively spliced at exon 3 that vary by tissue expression (21). The SLC25A3-B isoform is expressed ubiquitously, while the SLC25A3-A isoform is expressed primarily in striated muscle (22). Like other COX copper assembly factors, pathogenic variants in SLC25A3 lead to pathology that primarily presents as skeletal and cardiac myopathy (19, 23–26). Thus, understanding the role of SLC25A3 in mitochondrial copper transport in skeletal muscle is important to understanding the pathology of SLC25A3 dysfunction.

Skeletal muscle is a mitochondria-rich, highly organized, post-mitotic tissue that contains ~25% of total systemic copper and contributes to systemic metabolism (27–29). Muscle consists of multinucleated myofibers with closely associated stem cells that can be activated to regenerate skeletal muscle in the event of injury or disease. Once activated, skeletal muscle stem cells proliferate as myoblasts, differentiate, migrate, and then fuse to form a multinucleated myofiber. Activity of skeletal muscle stem cells is important for muscle regeneration and normal muscle homeostasis including maintenance of the neuromuscular junction and myonuclear homeostasis (30–34). Skeletal myoblasts derived from muscle stem cells exhibit varying metabolic requirements during differentiation including a consistent requirement for copper throughout the myogenic process. These requirements fluctuate between fatty acid oxidation (in quiescent muscle stem cells in vivo), glycolysis (myoblasts), and OXPHOS (myotubes) (35). However, the precise timing of when COX activity, and thus copper delivery to mitochondria, becomes essential is unknown.

Previous studies have shown that the *Slc25a3* gene is a target of the metal responsive transcription factor MTF1 that contributes to myoblast differentiation and that loss of SLC25A3 protein mildly impairs differentiation and reduces COX activity in either myoblasts or myotubes (36). Here, we investigated the importance of copper transport by SLC25A3 in differentiating myoblasts compared to fully differentiated myotubes. These studies reveal that SLC25A3 and COX assembly and activity are not required for myotube formation but are essential for survival of fully differentiated myotubes. We found that expression of either SLC25A3-A or -B isoform is sufficient to rescue myotube survival and that a variant of SLC25A3 that transports copper exclusively restores myotube survival to near wild type levels. Taken together, these data support a model wherein copper delivery to mitochondria is dispensable for myotube formation, but myotube survival depends on copper transport by SLC25A3 and COX activity. Thus, pathology related to SLC25A3 in skeletal muscle may be due to impaired copper delivery to mitochondria.

## Results

### SLC25A3 deficiency during myoblast differentiation leads to formation of small myotubes

A previous study showed that SLC25A3 deficiency led to decreased myoblast proliferation and reduced levels of differentiation markers when myoblasts were induced to differentiate, but the effect on SLC25A3 deficiency on myotube formation was not quantified directly (36). Here, siRNA was used to generate SLC25A3 deficiency in immortalized C2C12 and primary murine myoblasts prior to inducing them to differentiate with or without the extracellular copper chelator BCS (Figure 1A). Concentrations of BCS were chosen based on doses that do not inhibit myotube formation (37). Levels of proteins related to myoblast differentiation and copper metabolism were measured by immunoblot (Figure 1B). As expected, treatment with BCS led to increased levels of the copper chaperone for superoxide dismutase 1 (CCS), a protein that is stabilized when copper availability is low (38). The CuB containing COX subunit, COX2, did not exhibit a dramatic decrease in SLC25A3 deficient cells, and no change was detected in mitochondrial markers VDAC or TOMM20. Levels of the terminal differentiation marker embryonic myosin heavy chain (eMyHC) were reduced only when *Slc25a3* knockdown was combined with 50 µM BCS, suggesting that differentiating SLC25A3 deficient myoblasts may be sensitized to copper deprivation. Despite the reduced levels of eMyHC, *Slc25a3* knockdown did not inhibit myotube formation (Figure 1C). In fact, quantification of the fusion index (nuclei in multinucleated myotubes per total nuclei) revealed a slight increase in *Slc25a3* knockdowns in cells treated with 0 or 25 µM BCS (Figure 1D). However, SLC25A3 deficiency at 50 µM BCS led to the formation of smaller myotubes as quantified by myotube width (Figure 1E). Similar results were detected in *Slc25a3* knockdown primary myoblasts induced to differentiate with and without BCS (Figure 1F, Supplemental Figure 1A). Taken together, these data agree with existing literature and suggest that SLC25A3 deficiency may slow myoblast differentiation or inhibit secondary fusion or hypertrophy.

**Figure 1:**
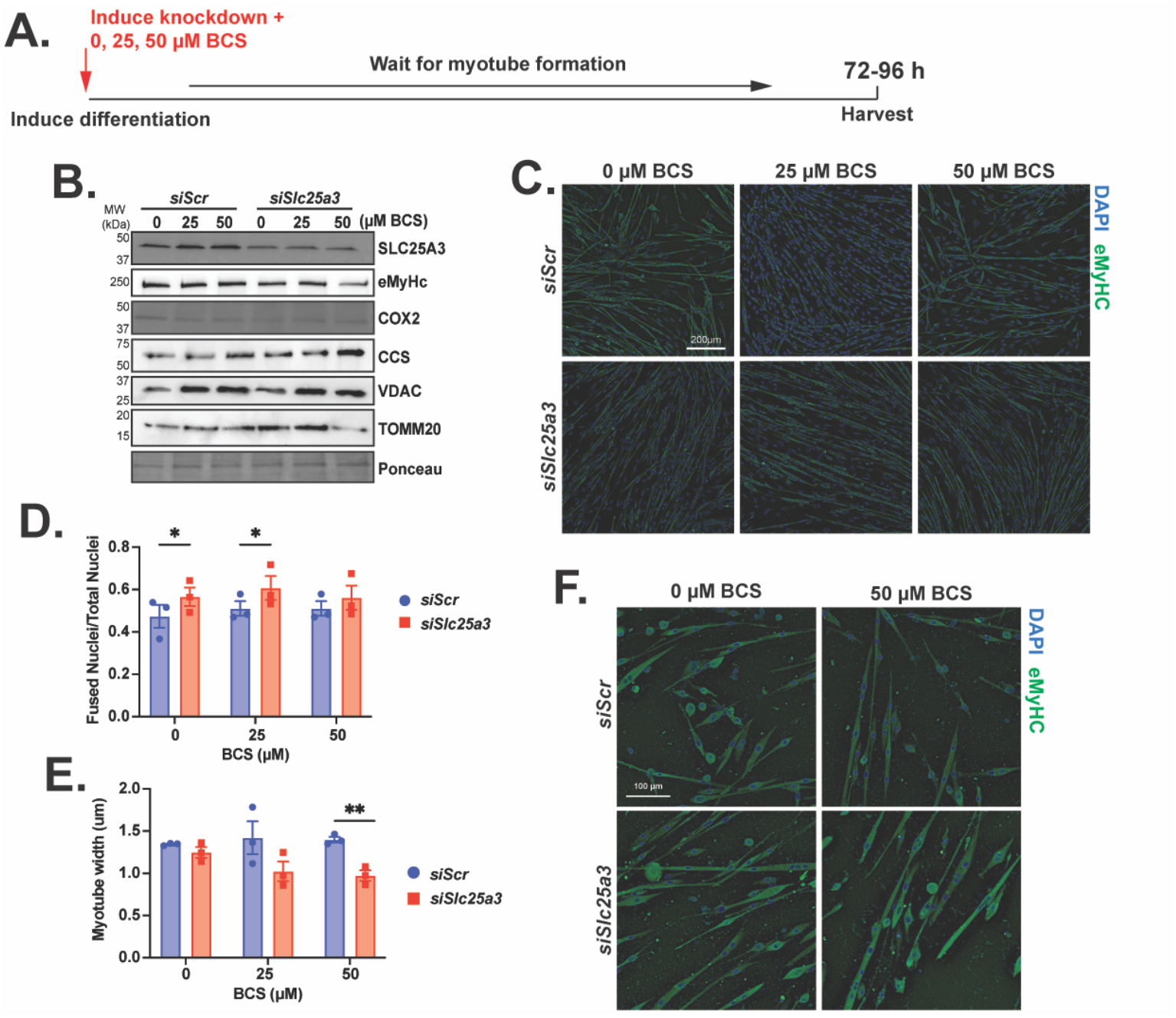
SLC25A3 deficiency during myoblast differentiation does not inhibit myotube formation. *A)* Schematic of experiment wherein siRNA was used to knock down *Slc25a3* at the same time that differentiation was induced. *B)* Immunoblot of C2C12 myotube lysates from myoblasts treated with negative control (*siScr*) or *Slc25a3* targeting siRNA (*siSlc25a3*) and differentiated in increasing concentrations of the copper chelator bathocuproine disulfonate (BCS). Blot was probed with antibodies to SLC25A3 to show knockdown, embryonic myosin heavy chain (eMyHC) as a differentiation marker, COX2 to show effect on CuA containing COX subunit, CCS as a measure of available copper, and VDAC and TOMM20 as controls for total mitochondrial content. Ponceau staining was used as a loading control. Blot is representative of n=3 biological replicates. *C)* Representative immunostained images of C2C12 myotubes from *siScr* and *siSlc25a3* myoblasts differentiated with or without BCS showing normal myotube formation across all conditions. Myotubes were stained with an antibody eMyHC and DAPI to visualize nuclei. Bar = 200 µm. *D)* Fusion index (ratio of nuclei in fused myotubes to total) in *siScr* and *siSlc25a3* myoblasts differentiated to myotubes with and without BCS showing significantly increased fusion index in *siSlc25a3* myotubes under low BCS conditions and no difference between *siScr* and *siSlc25a3* fusion index in 50 µM BCS. *E)* Myotube width in *siScr* and *siSlc25a3* myoblasts differentiated to myotubes with and without BCS showing significantly reduced width in *siSlc25a3* in 50 µM BCS. *F)* Representative images of primary murine myotubes differentiated with *siScr* or *siSlc25a3* with and without BCS. Bar = 100 µm. Additional replicates are shown in Supplementary Figure 1. For *D* and *E*, shown is mean ± SEM for n=3 biological replicates. Statistical significance was determined using two-way ANOVA with Sidak’s post-hoc testing. *p<0.05, **p<0.01.

### SLC25A3 deficiency leads to cell death in fully differentiated myotubes

Loss of function mutations in the human *SLC25A3* gene lead to skeletal and cardiac myopathy in children and adults but do not impair muscle development (39). To determine the effect of SLC25A3 deficiency in fully formed myotubes, an in vitro model of skeletal muscle, *Slc25a3* knockdown was induced for 48 hours after myotube formation was complete (Figure 2A). Loss of SLC25A3 in mature myotubes led to a marked decrease in detection of COX2 with little effect on TOMM20 (Figure 2B). As expected, levels of CCS increased in myotubes treated with BCS. Levels of eMyHC were unchanged. Surprisingly, SLC25A3 deficiency led to large-scale myotube detachment, suggesting cell death (Figure 2C). An ATP based cell viability assay indicated reduced myotube survival upon *Slc25a3* knockdown (Figure 2D). To determine if cell death precedes myotube detachment under SLC25A3 deficiency, viability was measured after only 24 hours of knockdown, before most cell detachment was observed (Figure 2E). Since SLC25A3 contributes to OXPHOS and ATP production, an alternative cell permeability-based assay was used. As expected, *Slc25a3* knockdown for 24 hours led to increased cell permeability, which is consistent with cell death (Figure 2F). Knockdown of *Slc25a3* in primary murine myotubes also led to cell death within 48 hours (Figure 2G, Supplemental Figure 1B). This result indicates that SLC25A3 supports survival of fully differentiated C2C12 and primary murine myotubes.

**Figure 2:**
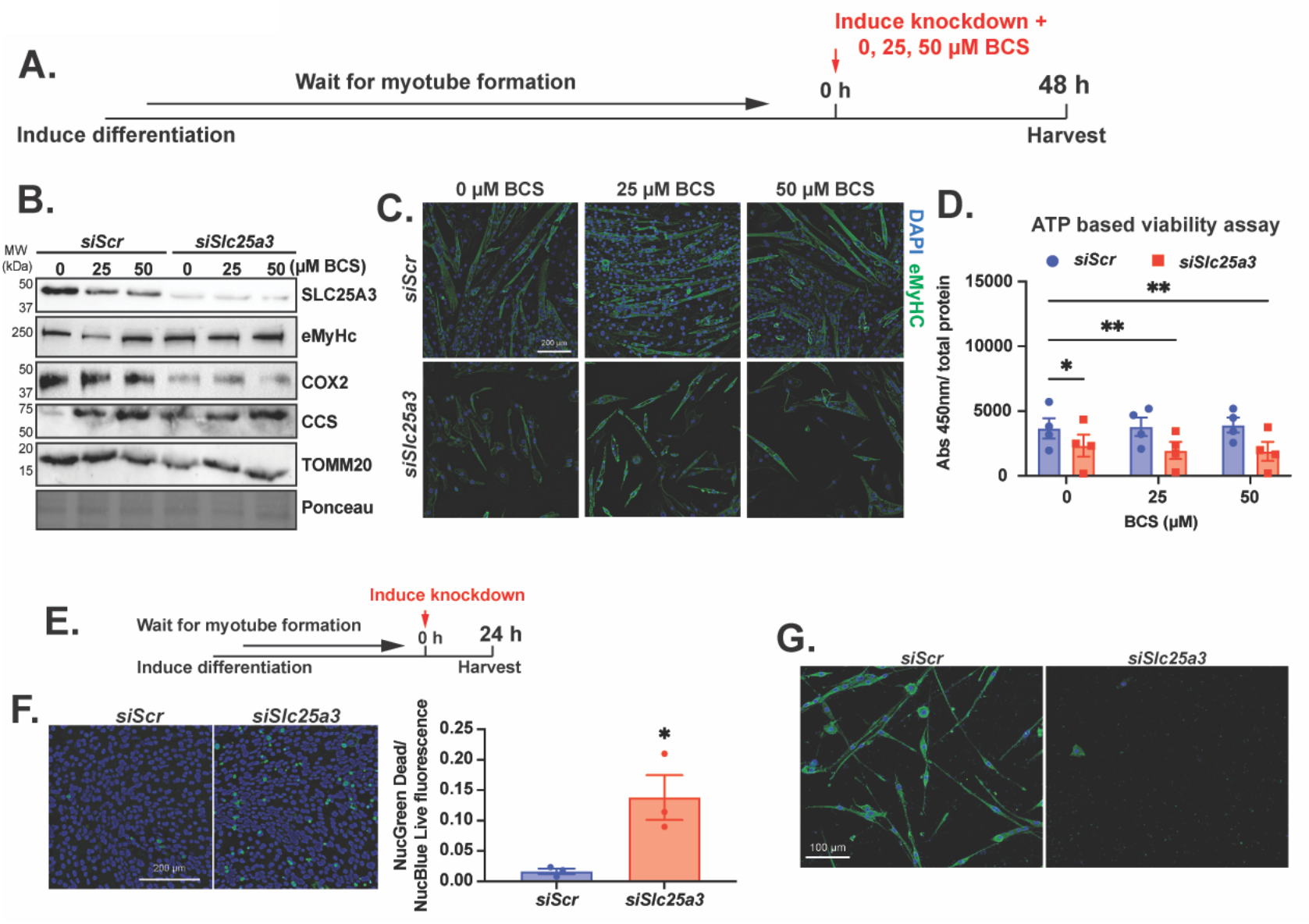
SLC25A3 deficiency in fully differentiated myotubes leads to cell death and detachment. *A)* Schematic of experiment wherein myoblasts were fully differentiated to myotubes and then siRNA used to knock down *Slc25a3* with or without BCS for 48 hours. *B)* Immunoblot of lysates from C2C12 myotubes treated with *siScr* and siSlc25a3 and increasing concentrations of BCS. Blots were probed with antibodies to SLC25A3 to show knockdown, embryonic myosin heavy chain (eMyHC) as a differentiation marker, COX2 to show effect on CuA containing COX subunit, CCS as a measure of available copper, and TOMM20 as a control for total mitochondrial content. Ponceau staining was used as a loading control. Blot is representative of n=3 biological replicates. *C)* Representative immunostained images of C2C12 myotubes from *siScr* and *siSlc25a3* myoblasts differentiated with or without BCS showing cell detachment and death in *siSlc25a3* myotubes. *D)* ATP based viability assay of C2C12 myotubes showing reduced viability in *siSlc25a3* in all BCS conditions. Shown is mean ± SEM for n=3 biological replicates. Statistical significance was determined using two-way ANOVA with Sidak’s post-hoc test. *p<0.05, **p<0.01. *E)* Schematic of short-term knockdown experiment wherein myoblasts were fully differentiated to myotubes and then siRNA used to knock down *Slc25a3* with or without BCS for 24 hours. *F)* Live/dead staining of C2C12 myotubes treated with siScr and *siSlc25a3*. Dead/permeable cells shown in green and nuclei (Hoechst staining) are shown in blue. Quantification revealed increased permeable/dead cells in *siSlc25a3* compared to *siScr* myotubes. Shown is mean ± SEM for n=3 biological replicates. Statistical significance was determined by one-tailed t-test. * p<0.05. *G)* Representative images of primary myotubes that were treated with *siScr* or *siSlc25a3* after differentiation was complete. Additional replicates are shown in Supplementary Figure 1.

The majority of pathogenic mutations in SLC25A3 are found in exon 3A, which encodes the striated muscle specific isoform SLC25A3-A (39). The alternatively spliced exons encoded by *Slc25a3-A* and *-B* isoforms differ little in their primary sequence, and the SLC25A3-A and -B proteins exhibit remarkably similar predicted structures and apparent affinity for copper transport but differ slightly in affinity for phosphate (Figure 3 A, B) (16, 40). Although the majority of striated muscle SLC25A3 is the A isoform, both isoforms are expressed (19). To determine whether isoform-specific differences contribute to myotube survival, sequence encoding either SLC25A3-A or -B was subcloned into the pcDNA 3.1 plasmid and expressed in fully differentiated myotubes concurrently with *Slc25a3* knockdown for 48 hours. No epitope tags were used to avoid disrupted transport by the tag, so expression of exogenous SLC25A3-A or -B was visualized as restoration of the SLC25A3 band by immunoblot (Figure 3C). The siRNA used in this experiment targets the *Slc25a3* 3’ untranslated region, so expression of exogenous *Slc25a3* coding sequence would not be expected to disrupt endogenous *Slc35a3* knockdown. Both SLC25A3-A and -B isoforms sufficiently rescued the myotube death phenotype (Figure 3D) as quantified by total green fluorescence representing eMyHC positive myotubes (Figure 3E). These data indicate that either SLC25A3-A or -B isoform is sufficient to promote myotube survival.

**Figure 3:**
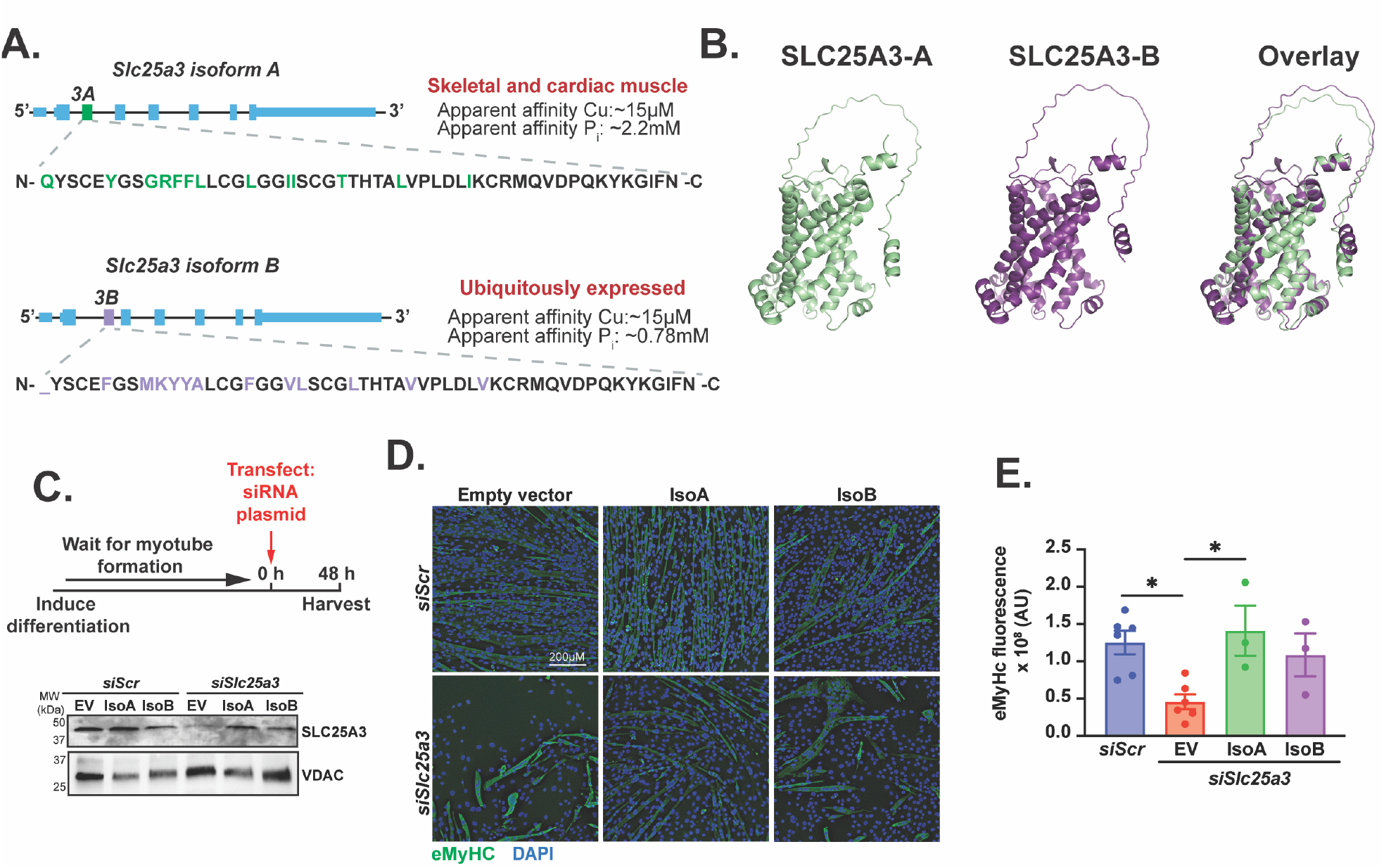
Both SLC25A3 A and B isoforms are sufficient to promote myotube viability. *A)* Schematic of *Slc25a3* RNAs showing alternatively spliced exons *3A* and 3B and corresponding protein sequence. *B)* Alpha-fold generated images of SLC25A3-A and -B and overlay showing - A and green and -B in purple. *C)* Schematic and immunoblot of rescue experiment where differentiated C2C12 myotubes were transfected with *siScr, siSlc25a3* with empty vector (EV) or vector encoding murine SLC25A3-A or -B. Blot was probed with an antibody to SLC25A3 to show knockdown and re-expression of -A and -B isoforms. An antibody to the mitochondrial voltage dependent anion channel (VDAC) was used as a loading control. Blot is representative of n=3 biological replicates. *D)* Immunofluorescence staining using an antibody to eMyHC showing restoration of myotube survival in *Slc25a3* knockdowns expressing exogenous SLC25A3-A or -B. *E)* Quantification of total eMyHC fluorescence showing significantly higher fluorescence in *siScr* or *siSlc25a3* plus SLC25A3-A compared to *siSlc25a3* plus EV. Shown is mean ± SEM for n=3 biological replicates. Statistical significance was determined using one-way ANOVA with Dunnett’s post-hoc test. *p<0.05.

### COX copper cofactor assembly and COX activity is required for myotube survival

As SLC25A3 contributes to Cu delivery to mitochondria for assembly into COX, we sought to better understand the COX requirements during myoblast differentiation. C2C12 myoblasts were induced to differentiate for 96 hours and harvested at different stages of differentiation (Figure 4A). Consistent with previous studies, Seahorse XR based assays indicated a shift in ATP production from glycolysis in myoblasts to OXPHOS in myotubes (Figure 4B) (35). As expected, levels of labile subunits from ETC complexes I-IV increased over myoblast differentiation along with total mitochondrial content as measured by VDAC (Figure 4C). Electron flow assays, where Seahorse XR based OCR is performed on live cells treated with individual ETC complex substrates and inhibitors, were used to measure the activities of Complexes I, II, and IV in proliferating myoblasts, myocytes in the early/intermediate stages of differentiation, and fully differentiated myotubes (Figure 4A) (41). While activity of Complex I rose substantially in myocytes and Complex II activity increased in myocytes before decreasing in myotubes, activity of COX/Complex IV increased in myocytes and continued to increase in myotubes (Figure 4D), suggesting that COX activity does not reach maximal levels until differentiation is complete. This result is consistent with a requirement for copper delivery to COX in fully differentiated myotubes.

**Figure 4:**
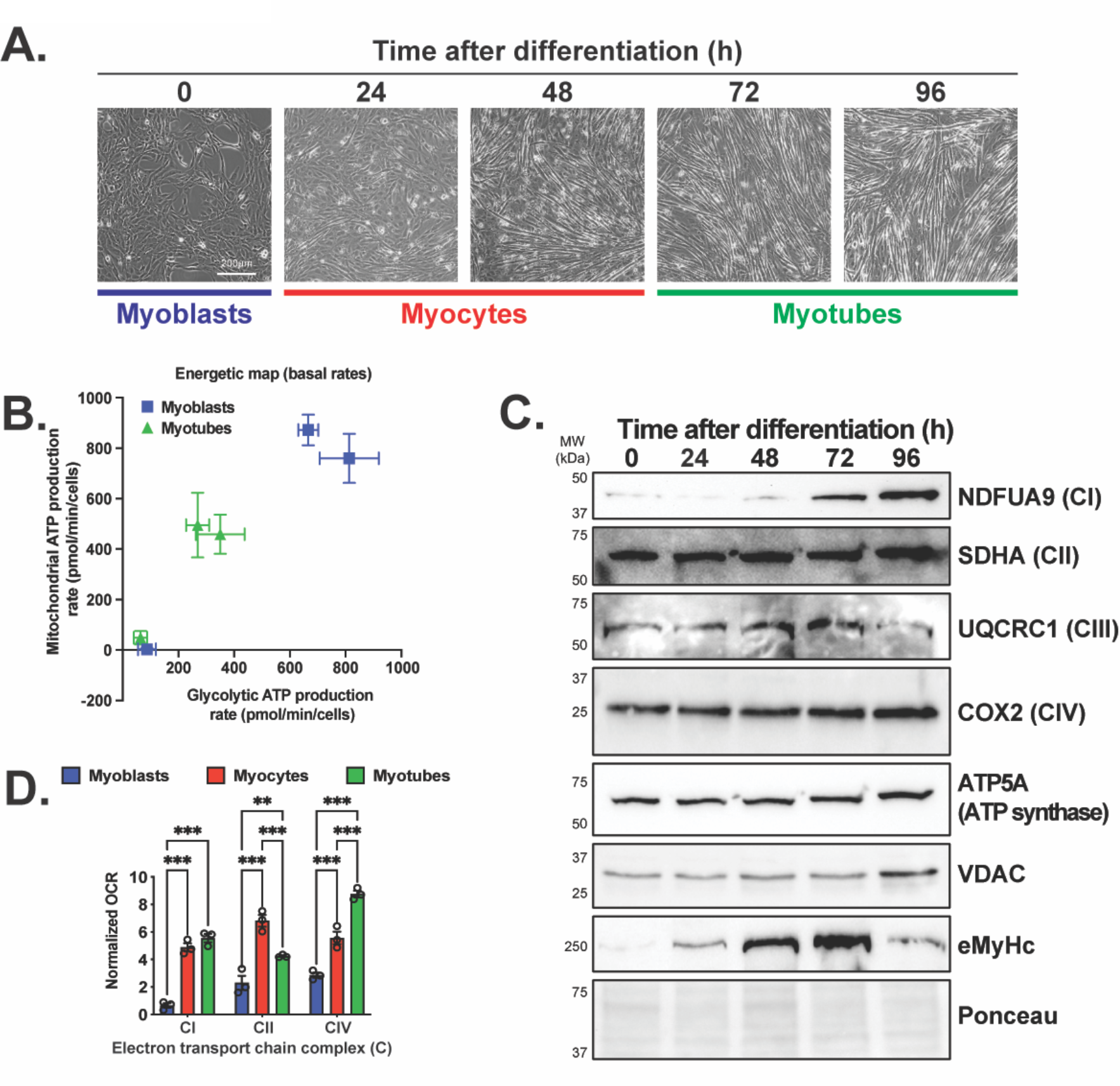
Myoblast differentiation is associated with increase in ATP generation from OXPHOS. *A)* Representative images showing different stages of C2C12 myoblast differentiation over time. *B)* Seahorse-based assay of oxygen consumption versus extracellular acidification showing increased rate of mitochondrial (y axis) versus glycolytic (x axis) ATP production in differentiated myotubes versus myoblasts. Shown are individual data points for three separate biological replicates. *C)* Immunoblot showing increased levels of labile subunits of the electron transport chain complexes (CI-CIV) and ATP synthase at different timepoints during myoblast differentiation. eMyHC was used as a marker of differentiation and Ponceau stain was used as a loading control. *D)* Seahorse-based electron flux assay showing early elevation in activities of complex I (CI) in myocytes, an increase in succinate dehydrogenase/complex II (CII) in myocytes that decreases in myotubes, and a steady increase in COX/complex IV (CIV) that reaches its maximum only after differentiation is complete. Shown is mean ± SEM for n=3 biological replicates. Statistical significance was determined using two-way ANOVA with Sidak’s post-hoc test. ***p<0.001.

To confirm the importance of copper delivery to COX in myoblast differentiation and myotube survival, we investigated the roles of various factors related to COX enzyme complex assembly and copper cofactor delivery (Figure 5A). The majority of copper center assembly factors for COX are localized to the IMS (4). The membrane tethered SCO1 and SCO2 proteins along with COA6 contribute to copper delivery to the Cu_A_ site in COX2 while COX19 and COX11 provide copper to the Cu_B_ site in COX1. Upstream, the soluble IMS protein COX17 delivers copper to the other assembly factors. While not directly implicated in copper delivery to COX, the translational activator for COX1 (TACO1) binds to the mitochondrial ribosome to prevent stalling at polyproline tracts in COX1 and promotes its translation (42, 43). As expected, steady-state RNA and protein levels of COX assembly factors increased as myoblast differentiation proceeded (Figure 5B, C). To directly impair copper delivery to COX, siRNA was used to knock down COX17 in myoblasts prior to inducing differentiation in cells with or without BCS treatment. Treatment with BCS led to increased levels of CCS as expected, and *Cox17* knockdown led to a marked decrease in the levels of eMyHC (Figure 5D). A slight decrease in COX2 was detected, which was exacerbated by the addition of BCS and no change in total mitochondrial content was detected as measured by levels of VDAC and TOMM20. Similar to the result observed in SLC25A3 deficiency, myotube formation itself was not inhibited by *Cox17* knockdown (Figure 5E). However, COX17 deficiency during myoblast differentiation led to reduced myotube width even without BCS treatment (Figure 5F). When COX17 deficiency was induced by siRNA in fully differentiated myotubes with and without BCS, CCS protein was increased in myotubes treated with BCS, levels of COX2 decreased, and no change was detected in eMyHC, VDAC, or TOMM20 (Figure 5G). Like SLC25A3 deficiency, COX17 deficiency led to detachment and death of fully differentiated myotubes as quantified by total eMyHC fluorescence (Figure 5H, I). These indicate copper delivery to COX is required for myotube survival but not formation.

**Figure 5:**
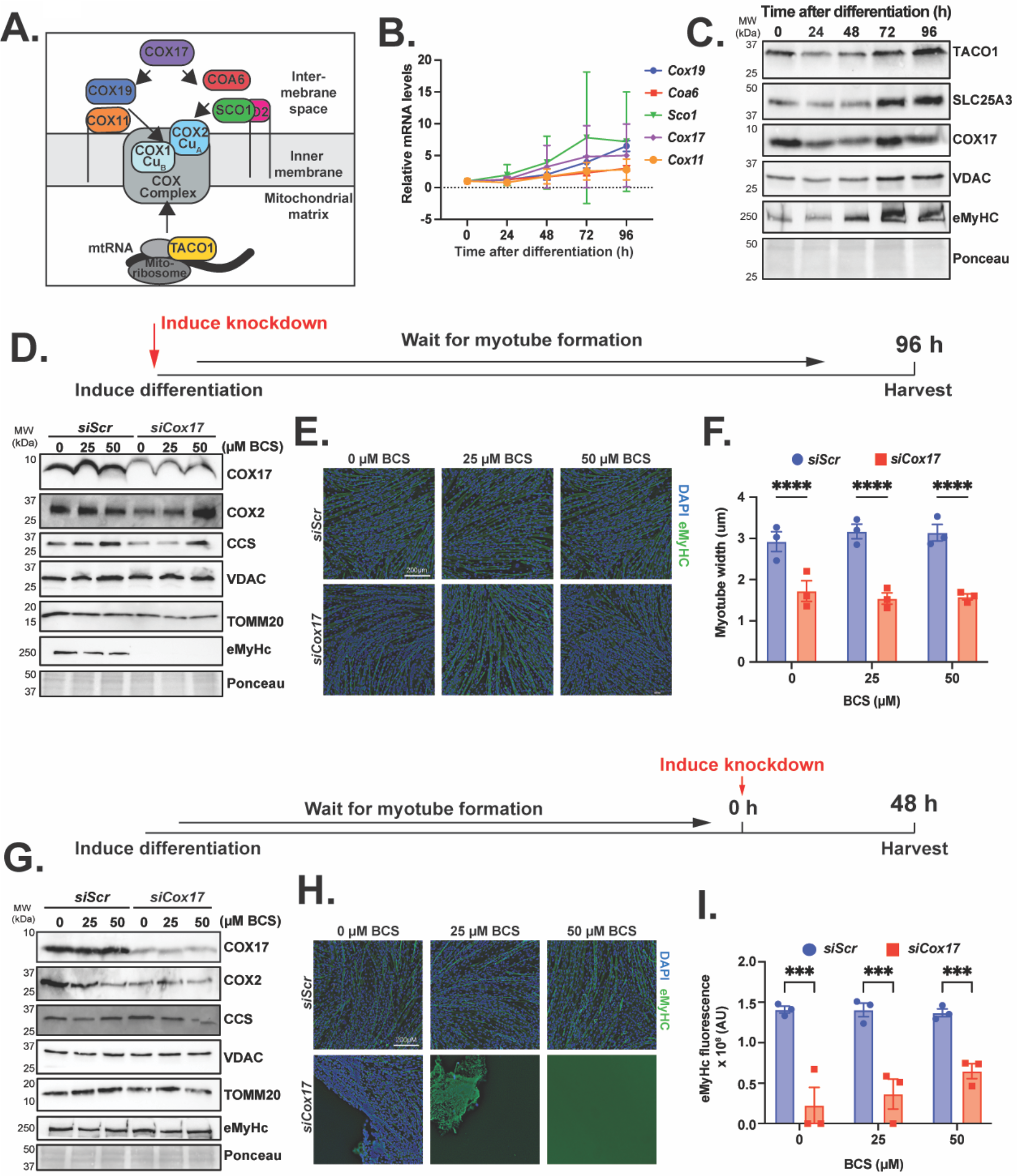
Deficiency of COX Cu cofactor assembly factor COX17 does not impair myotube formation but limits myotube viability. *A)* Schematic of COX assembly factors in the mitochondrial matrix and intermembrane space. *B)* Quantitative PCR showing increasing levels of transcripts encoding COX copper cofactor copper chaperones during myoblast differentiation. Shown is mean ± SEM for n=3 biological replicates. *C)* Immunoblot showing increasing levels of TACO1, SLC25A3, and COX17 during differentiation. VDAC was used as a mitochondrial marker showing increased mitochondrial content during myoblast differentiation. Ponceau stain was used as a loading control. Blot is representative of n=3 biological replicates. *D)* Schematic and immunoblot showing COX17 deficiency induced by *Cox17* targeting siRNA (siCox17) with or without BCS immediately prior to inducing differentiation. Immunoblot shows lysates probed with antibodies to COX17 to show knockdown, eMyHC as a differentiation marker, COX2 to show effect on CuA containing COX subunit, CCS as a measure of available copper, and TOMM20 as a control for total mitochondrial content. Ponceau staining was used as a loading control. Blot is representative of n=3 biological replicates. *E)* Representative immunostained images of C2C12 myotubes from *siScr* and *siCox17* myoblasts differentiated with or without BCS showing normal myotube formation across all conditions. Myotubes were stained with an antibody eMyHC and DAPI to visualize nuclei. Bar = 200 µm. *F)* Myotube width in *siScr* and *siCox17* myoblasts differentiated to myotubes with and without BCS showing significantly reduced width in *siCox17* in 50 µM BCS. *G)* Schematic and immunoblot showing COX17 deficiency induced by *Cox17* targeting siRNA (*siCox17*) with or without BCS in fully differentiated myotubes. Immunoblot was probed with antibodies listed in *D*. Blot is representative of n=3 biological replicates. *H)* Representative immunostained images of C2C12 myotubes from *siScr* and *siCox17* myoblasts differentiated with or without BCS showing cell detachment and death in *siCox17* myotubes. *I)* Quantification of total eMyHC fluorescence showing significantly lower fluorescence in *siCox17* compared to *siScr*. For *F* and *I*, shown is mean ± SEM for n=3 biological replicates. Statistical significance was determined using two-way ANOVA with Sidak’s post-hoc test. ***p<0.001, ****p<0.0001.

We then targeted COX assembly and activity directly to confirm the COX requirement in myotube formation versus viability. Knockdown of *Taco1* prior to inducing myoblast differentiation (Figure 6A) failed to inhibit myotube formation, though addition of the BCS slightly reduced myotube size (Figure 6B). As expected, levels of CCS increased upon BCS treatment but were not affected by *Taco1* knockdown (Figure 6A). The *Taco1* siRNA was not sufficient to knock down TACO1 in fully differentiated myotubes (data not shown), which is likely due to the stability of the TACO1 protein. Instead, we inhibited COX activity directly using potassium cyanide (KCN) in C2C12 and primary murine myoblasts during differentiation (Figure 6C) (44). COX inhibition with KCN did not impede myotube formation (Figure 6D). However, as was the case with SLC25A3 or COX17 deficiency, myotube width was reduced (Figure 6E). The fusion index in primary myoblasts treated with KCN was not affected (Figure 6F). Chemical inhibition of ETC complexes II or III, and ATP synthase yielded similar results where myotube formation was not impaired but fully differentiated myotubes exhibited reduced survival (Supplemental Figure 2), supporting a model wherein ETC activity is dispensable for myoblast differentiation but required for myotube survival.

**Figure 6:**
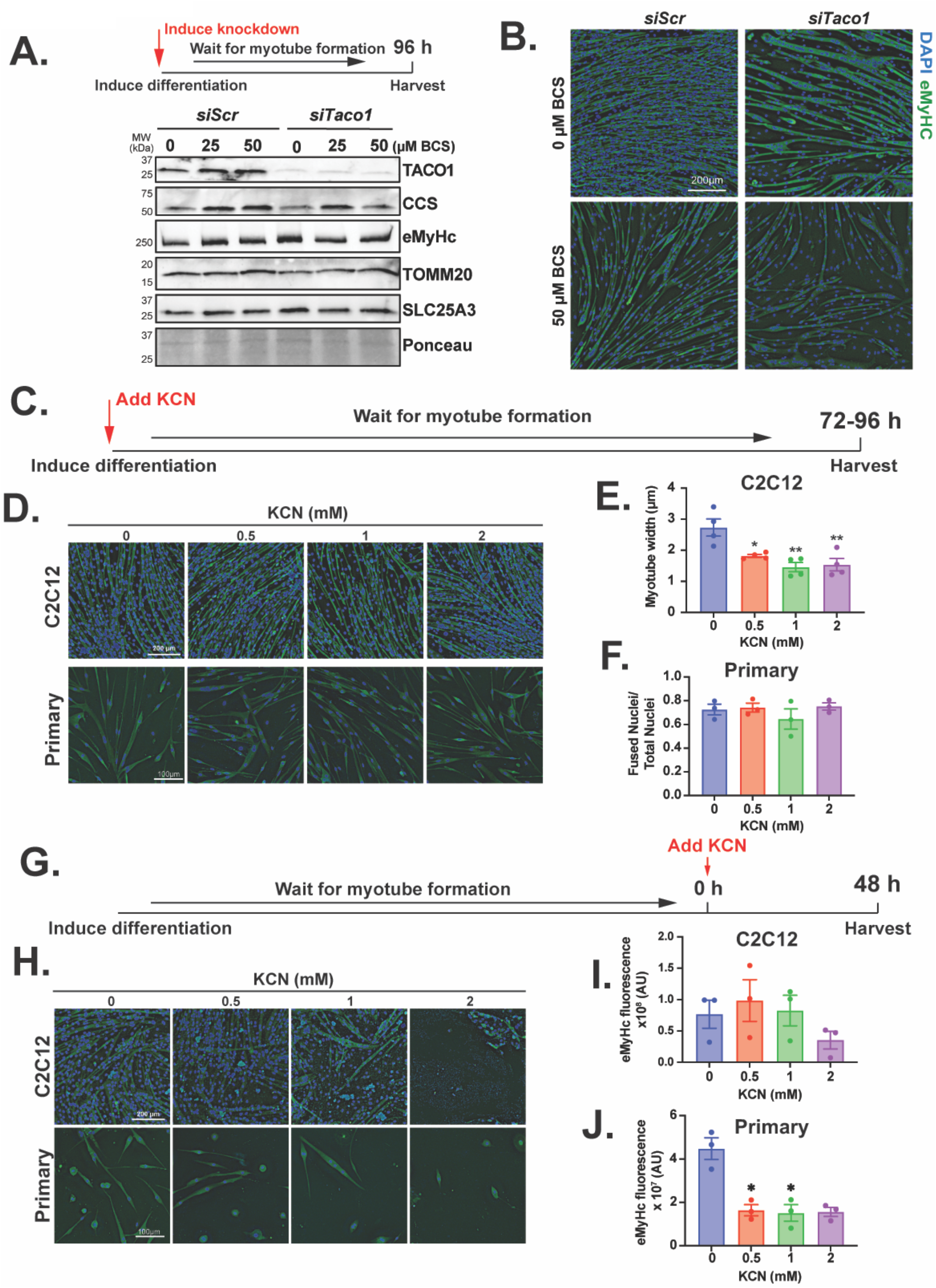
Chemical inhibition of COX activity does not prevent myotube formation but impairs myotube viability. *A)* Schematic of experiment and immunoblot showing TACO1 knockdown induced by siRNA (*siTaco1*) in myoblasts prior to inducing differentiation to myotubes with or without addition of BCS. Immunoblot shows lysates probed with antibodies to TACO1 to show knockdown, CCS as a measure of available copper, eMyHC as a differentiation marker. An antibody to TOMM20 was used a control for total mitochondrial content. Ponceau staining was used as a loading control. Blot is representative of n=3 biological replicates. *B)* Representative immunostained images of C2C12 myotubes from *siScr* and *siTaco1* myoblasts differentiated with or without BCS showing normal myotube formation across all conditions. Myotubes were stained with an antibody eMyHC and DAPI to visualize nuclei. Images are representative of n=3 biological replicates. Bar = 200 µm. *C)* Schematic of experiment wherein COX activity was inhibited with potassium cyanide (KCN) as myoblast differentiation was induced. *D)* Representative immunostained images showing normal myotube formation in C2C12 (top) and primary (bottom) myoblasts induced to differentiate with variable concentrations of KCN. *E)* Quantification of myotube width showing reduced myotube size when C2C12 myoblasts were differentiated in the presence of KCN. Myotubes were stained with an antibody eMyHC and DAPI to visualize nuclei. Bar = 200 µm for C2C12 myotubes and 100 µm for primary myotubes. *F)* Fusion index showing no impaired myotube formation in primary myoblasts induced to differentiate in the presence of KCN. *G)* Schematic of experiment wherein myoblasts were fully differentiated to myotubes and then treated with KCN for 48 hours. *H)* Representative immunostained images showing fully differentiated C2C12 (top) and primary (bottom) myotubes treated with variable doses of KCN. Myotubes were stained with an antibody eMyHC and DAPI to visualize nuclei. Bar = 200 µm for C2C12 and 100 µm for primary myotubes. *I)* Quantification of eMyHC fluorescence in C2C12 myotubes and *J)* primary myotubes showing reduced fluorescence associated with cell detachment and death. For E, F, I, and J, shown is mean ± SEM for n=3-4 biological replicates. Statistical significance was determined using one-way ANOVA with Dunnett’s post-hoc test. *p<0.05, **p<0.01.

### Copper transport activity by SLC25A3 is important for myotube survival

Although SLC25A3 is known to transport both copper and phosphate, it is not known how each of these transport activities contribute to skeletal muscle. To confirm that copper is required for myotube survival, fully differentiated myotubes were treated with the intracellular copper chelator tetrathiomolybdate (TTM). Copper chelation by TTM led to detachment of fully differentiated myotubes (Supplemental Figure 3), indicating that copper is required for myotube survival. To determine if the myotube death phenotype caused by SLC25A3 deficiency is related to mitochondrial copper deficiency, we knocked down *Slc25a3* in myotubes in the presence of exogenous copper sulfate (CuSO_4_) (Figure 7A). Addition of 50 or 100 µM CuSO_4_ rescued cell death in *Slc25a3* knockdown myotubes (Figure 7B) as quantified by total eMyHC fluorescence (Figure 7C). A recent study reported that a single alanine substitution at leucine 175 in the SLC25A3-B isoform permits transport of copper but not phosphate in mouse embryonic fibroblasts (MEFs) (45). Here, we generated a single alanine substitution at leucine 176, the equivalent position in the SLC25A3-A isoform, to explore the importance of copper transport activity in myotube survival (Figure 7D). We also generated an alanine variant at histidine 76, which disrupts both copper and phosphate transport by SLC25A3. Expression of both variants was visualized as restoration of SLC25A3 detection by immunoblot (Figure 7E). Myotube survival as visualized by immunofluorescence staining using an antibody to eMyHC (Figure 7F) revealed that expression of the SLC25A3-A Leu176Ala variant rescued myotube survival while expression of the His76Ala did not. Quantification of total eMyHC fluorescence indicated that *Slc25a3* knockdown with empty vector or expressing the His76Ala variant showed significantly lower eMyHC fluorescence than scrambled control while the Leu176Ala variant was restored to near control levels (Figure 7G), indicating that the myotube death phenotype was reversed by expressing a variant of SLC25A3-A that can transport copper only. Taken together, these data demonstrate that copper transport by SLC25A3 is critical for myotube survival.

**Figure 7:**
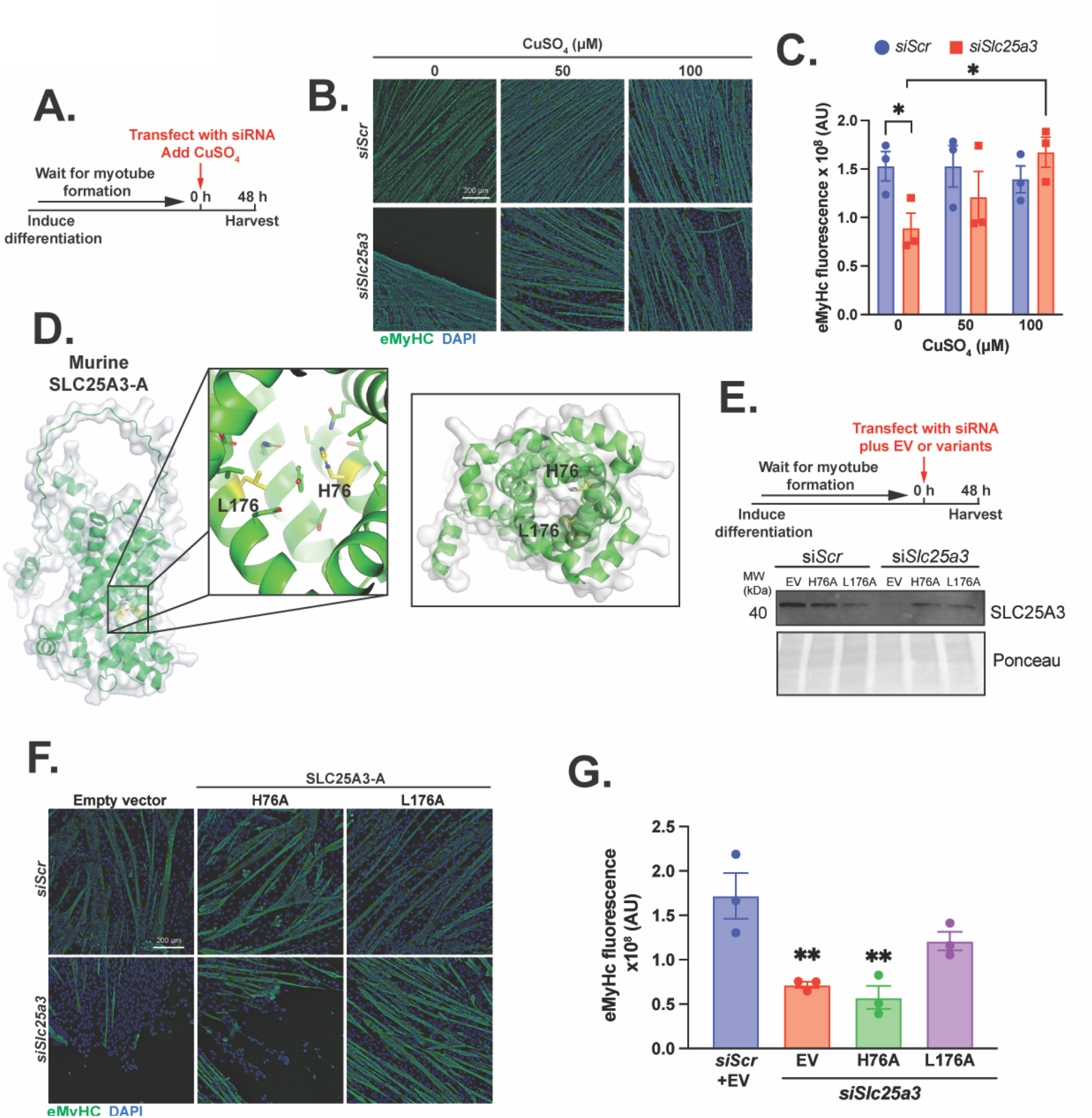
Exogenous copper or expression of a SLC25A3 that exclusively transports copper is sufficient to myotube viability in *Slc25a3* deficient cells. *A)* Schematic of experiment wherein *Slc25a3* was knocked down in fully differentiated myotubes with or without added exogenous copper. *B)* Representative immunostained images of *siScr* and *siSlc25a3* myotubes with or without added copper (CuSO_4_) showing cell detachment and death in non-treated but not copper treated *siSlc25a3* myotubes. Myotubes were stained with an antibody eMyHC and DAPI to visualize nuclei. Bar = 200 µm. *C)* Quantification of total eMyHC fluorescence showing significantly lower fluorescence in non-treated *siSlc25a3* myotubes compared to *siScr* or *siSlc25a3* myotubes treated with CuSO_4_. Shown is mean ± SEM for n=3 biological replicates. Statistical significance was determined using two-way ANOVA with Sidak’s post-hoc test. *p<0.05. *D)* Alpha-fold model of murine SLC25A3 isoform A (SLC25A3-A) highlighting residues His76 (H76) and Leu176 (L176) that are essential for the general transport mechanism or phosphate transport by SLC25A3. *E)* Schematic of experiment wherein myotubes were transfected with siSlc25a3 with empty vector (EV) or vector encoding H76A or L176A and immunoblot of lysates probed with an antibody to SLC25A3 to show knockdown and expression of variants. Ponceau staining was used as a loading control. Blot is representative of n=3 biological replicates. *F)* Representative immunostained images of *siScr* and *siSlc25a3* myotubes with or without expression of SLC25A3 H76A or L176A variants. Myotubes were stained with an antibody eMyHC and DAPI to visualize nuclei. Bar = 200 µm. *G)* Quantification of total eMyHC fluorescence showing significantly lower fluorescence in *siSlc25a3* transfected with empty vector (EV) or H76A variant compared to *siScr*. Shown is mean ± SEM for n=3 biological replicates. Statistical significance was determined using one-way ANOVA with Dunnett’s post-hoc test. **p<0.01.

## Discussion

Here, we aimed to understand the importance of copper delivery to mitochondria during skeletal myoblast differentiation. We showed that deficiency of the dual copper and phosphate carrier SLC25A3 impairs but does not inhibit myotube formation. Instead, loss of SLC25A3 leads to a pronounced cell death phenotype when induced in fully differentiated myotubes. This result was replicated using *Cox17* knockdown to impair COX copper cofactor assembly and by direct inhibition of COX or other complexes in the ETC. These phenotypes are consistent with the well-documented metabolic shift from glycolysis to OXPHOS as myoblasts differentiate, which correlates with a late increase in COX activity as identified here using electron flow assay. Expression of either SLC25A3-A or -B isoform was sufficient to promote survival in *Slc25a3* knockdown myotubes. Importantly, survival of SLC25A3 deficient myotubes was restored by exogenous copper or expression of an SLC25A3 variant that transports copper but not phosphate. Taken together, the data presented here highlight the importance of copper transport by SLC25A3 in skeletal muscle tissue and indicate that copper delivery to mitochondria for COX is critical for survival of myotubes but dispensable for their formation.

Several studies have established the importance of SLC25A3 in both copper and phosphate transport to mitochondria, but few have interrogated the importance of each substrate for mitochondrial function (16, 20, 40, 45–47). A detailed evolutionary history of the *Saccharomyces cerevisiae* homolog of SLC25A3, Pic2, alongside variant screening of conserved residues in Pic2 and SLC25A3 was recently reported (45). Mutation of a highly conserved leucine residue, Leu 175 in SLC25A3-B, was able to restore copper but not phosphate transport in SLC25A3 knockout MEFs. A more recent study showed rescue of mitochondrial copper and mitochondrial flickering by the SLC25A3-B Leu175Ala variant in an OPA1 knockout cell line (47). For this study, we generated an alanine substitution at the corresponding SLC25A3-A leucine 176 and showed nearly complete rescue of the cell death phenotype in *Slc25a3* knockdown myotubes, indicating that it is the copper transporting activity of SLC25A3 that is required for myotube survival. These results agree with a published study in primary myoblast differentiation showing that SLC25A3 can co-precipitate with copper-containing subunits of COX and the COX copper assembly protein SCO1 and highlight the importance of copper transport by SLC25A3 in skeletal muscle cells (36).

The relative contribution of copper versus phosphate transport by SLC25A3 is not known. Other members of the mitochondrial carrier family likely contribute to transport of both substrates. For example, the *S. cerevisiae* mitochondrial iron carrier Mrs3 contributes to copper import as Mrs3 can mediate copper transport in vitro and simultaneous deletion of *mrs3* and *pic2* genes cause a copper dependent respiratory growth defect in yeast (48). Whether the mammalian Mrs3 homologs, mitoferrin1 (SLC25A37) and mitoferrin 2 (SLC25A28), contribute to copper import remains unexplored. The ATP-Mg/phosphate (APC) carriers (SLC25A23, SLC25A24, SLC25A25) have been proposed to use phosphate antiport as a mechanism for importing adenine nucleotides into the mitochondrial matrix (17). In vitro studies showed that the APC carriers are capable of transporting phosphate or adenine nucleotides in either direction, so they may contribute to phosphate import into the matrix in some contexts (49). Similarly, the two SLC25A3 isoforms may exhibit differential contributions to copper and phosphate transport in skeletal muscle. Additional studies are needed to understand the details of copper versus phosphate transport activity by SLC25A3 and other mitochondrial carrier family proteins in striated muscle.

Unlike other diseases caused by impaired COX assembly, symptoms caused by loss of SLC25A3 are largely confined to skeletal and cardiac muscle due to the majority of pathogenic variants being found within the striated muscle specific *SLC25A3-A* isoform (39). Though SLC25A3-A is expressed only in muscle, the SLC25A3-B isoform is expressed ubiquitously, including in muscle. Here, myotube viability caused by SLC25A3 deficiency was rescued by expressing either SLC25A3-A or -B isoform, suggesting that some critical functions can be mediated by SLC25A3-B. Given the potential for SLC25A3-B to reverse the phenotype of total *Slc25a3* knockdown, perhaps raising levels of SLC25A3-B might relieve symptoms caused by loss of SLC25A3-A in striated muscle. Alternatively, copper supplementation provided to individuals with SLC25A3-A myopathy may relieve skeletal muscle weakness. More studies of the SLC25A3 isoforms, including the mechanisms mediating mutually exclusive splicing of exons 3A and 3B, are needed.

SLC25A3 deficiency in murine cardiac muscle leads to widespread alterations in mitochondria including mitochondrial fragmentation and cristae disorganization (20, 50). Treatment of heart-specific *Slc25a3* knockout mice with meclizine, an antihistamine that inhibits OXPHOS and promotes glycolysis, can improve cardiac muscle function in part by raising levels of the MICOS complex that controls organization of cristae junctions (51). These results raise the possibility that SLC25A3 may play some important structural role in mitochondrial organization. However, we report here that a His76Ala variant, which ablates transport of either copper or phosphate by SLC25A3, failed to rescue the cell death phenotype caused by SLC25A3 deficiency in skeletal myotubes. Thus, it is likely that transport activity by SLC25A3 is essential in skeletal muscle.

It is difficult to compare the myoblast differentiation and myotube death phenotypes reported here to the existing literature, though overall our data agree with other studies reporting differential requirements for mitochondrial activity depending on myoblast differentiation state. Most recently a study by Arnold and colleagues reported that proliferating myoblasts employ a non-canonical TCA cycle leading to accumulation of cytosolic citrate and a shift to the canonical TCA cycle upon differentiation (52). Though the importance of mitochondrial biogenesis in myoblast differentiation is well known, few studies have shown the effect of direct ETC inhibition on myotube formation (35). One study showed that inhibition of complexes I or II impaired myotube formation in culture while inhibition of complexes III or IV did not, though the concentrations used were much lower than those used here (53). However, a recent report of reverse electron transfer in a model of facioscapulohumeral muscular dystrophy (FSHD) showed that the complex II inhibitor dimethyl malonate did not inhibit formation of myotubes formed from control myoblasts (54). Interestingly, the complex I inhibitor rotenone does impair myotube formation, though this is through an off-target effect on the Raf1/ROCK signaling cascade (55). Similarly, the mitochondrial ribosome inhibitor chloramphenicol inhibits myotube formation, but this could be due to the fact that rotenone impairs mitochondrial biogenesis (56). Short-term inhibition of complex I with piericidin in myoblasts led arginine deficiency and activation of the integrated stress response (ISR) while in myotubes inhibiting ATP synthase led to inner membrane hyperpolarization and ISR activation (57). However, this study used only 10 hours of inhibitor treatment and complex IV was not targeted. To our knowledge, no comparative studies directly inhibiting complex IV during myoblast differentiation and in myotubes have been performed. Muscle stem cell specific knockout of murine *Cox10*, which encodes a farnesyltransferase critical for the COX heme *a* cofactor biosynthesis, led to reduced COX activity and aberrant muscle stem cell fusion leading to exhaustion, but an effect on myoblast differentiation was not measured directly (58). In mature murine skeletal muscle, COX10 knockout led to severe muscle wasting, which is distinct from the in vitro cell death phenotype shown here caused by COX inhibition or disrupted copper delivery to COX (59). This discrepancy could be due to the inherent differences between cultured C2C12 or primary myotubes and skeletal muscle in vivo as COX10 knockout in skeletal muscle led to systemic metabolic shifts that may represent an adapted muscle niche (60). Though cardiac muscle specific knockout of *Slc25a3* has been performed in mice, the corresponding skeletal muscle studies are needed to better understand the skeletal muscle pathology of SLC25A3 deficiency.

In this study we showed that copper transport by SLC25A3 is important to support survival in skeletal myotubes, a cell culture model analogous to a major tissue affected by pathologic SLC25A3 deficiency. This study emphasizes the importance of copper delivery to mitochondria in skeletal muscle but raises additional mechanistic questions. While SLC25A3 is instrumental in delivering copper to the mitochondrial matrix, how this copper is routed from the matrix to the intermembrane space for coordinated assembly into COX is not known. Similarly, little is known about how copper is trafficked from the plasma membrane to mitochondria. The COX17 copper chaperone is dual localized to the cytosol and mitochondrial intermembrane space and has been proposed to carry copper to mitochondria (61). Indeed, in some studies, COX17 deficiency reduces total mitochondrial copper (62, 63). However, in *S. cerevisiae*, Cox17 tethered to the mitochondrial inner membrane is sufficient to ensure adequate copper delivery to COX, suggesting that an alternate pathway carries copper to mitochondria (64). A recent study revealed differential changes in the proteomes of neuronal cells deficient in either COX17, which leads to reduced mitochondrial copper, and the plasma membrane copper importer CTR1, which leads to reduced mitochondrial and cytoplasmic copper (62). The results of this study suggest that COX17 may contribute to other cellular functions or contribute to copper homeostasis in other ways beyond providing copper to COX. Another recent study implicated the outer membrane mitochondrial carrier family protein MTCH2 (SLC25A50) in modulating mitochondrial copper in skeletal muscle, though the molecular details remain to be determined (65). Finally, the mitochondrial copper pool appears capable of driving intracellular and systemic signaling, suggesting that the mitochondrial pool is critical for regulating overall copper homeostasis (5, 66–68). Considering that skeletal muscle represents a major regulator of systemic metabolism, the role of mitochondrial copper in skeletal muscle as it contributes to systemic metabolism should be explored (69). The sum of the data presented here and in these recent studies indicate that copper delivery to mitochondria is critical in skeletal muscle, but more work is needed to elucidate the finer mechanistic details and broader impacts of this pathway.

## Experimental procedures

### Cell culture

C2C12 cells were purchased from American Type Culture Collection (ATCC) and regularly validated based on ability to form myotubes. C2C12 were grown in Dulbecco’s modified Eagle’s medium (DMEM) containing 4.5 g/L glucose and supplemented with L-glutamine and sodium pyruvate (Corning MT10013CV) in the presence of 10% fetal bovine serum (FBS) (Cytiva SH3091003), 50 µg/ml penicillin and streptomycin (pen/strep) (Corning MT30001CI), and 2.5 µg/ml Plasmocin (Invitrogen, NC9698402) at 37°C in 5% CO_2_. To induce C2C12 myoblast differentiation, cells were grown to 80-90% confluence and medium was switched to DMEM with 2% horse serum (Cytiva, SH3007403) and 50 µg/ml pen/strep for approximately 96 hours. For experiments using myocytes, differentiating C2C12 cells were harvested when elongated myocytes were visible, approximately 48 hours after inducing differentiation. Myotubes were treated or harvested for all samples when elongated myotubes were visible in control samples, approximately 72-96 hours after inducing differentiation.

Primary myoblasts isolated from wild type adult C57Bl/6J murine limb muscle were thawed from existing cell stocks and grown as previously described (70) on tissue culture dishes coated with collagen I (Corning CB-40231) in Ham’s F10 medium (Gibco 11550043) containing 20% FBS, 50 µg/ml pen/strep, and 2.5 µg/ml plasmocin supplemented with 5 ng/ml human basic fibroblast growth factor (Pepro Tech 100-18B). Primary myoblasts were regularly validated based on morphology (small, round) and ability to form myotubes. Cells were differentiated on plates coated with enactin-collagen IV-laminin (Sigma 08-110) in DMEM containing 1 g/L glucose supplemented with L-glutamine and sodium pyruvate (Gibco 11885084) with 50 µg/ml pen/strep and 1% insulin-selenium-transferrin solution (Gibco 51300044). Differentiation was considered complete when myotube formation was visible in control samples, approximately 72 hours after inducing differentiation.

Treatment of differentiating myoblasts or fully differentiated myotubes with copper chelators or ETC inhibitors was performed using concentrations based on previous studies (37, 44, 53, 71–74). Bathocuproine disulfonate (BCS, Sigma B1125) or fresh ammonium tetrathiomolybdate (TTM, Sigma 323446) to chelate copper, or exogenous copper provided as copper sulfate (CuSO_4_, Sigma 209198) were suspended in water and sterile filtered using a 0.22 µm syringe filter prior to adding to differentiation medium. Inhibitors of ETC complexes II (oxaloacetate, ThermoFisher AA1578909) and IV (potassium cyanide, Sigma 207810) were resuspended in water and inhibitors for complex III (antimycin A, ThermoFisher AAJ63522LB0) and ATP synthase (oligomycin, Sigma 495455) were resuspended in DMSO. All chemical reagents resuspended in water were filtered using a 0.22 µm syringe filter.

### Genetic knockdown induced with dicer substrate siRNAs

Deficiency of *Slc25a3, Cox17*, or *Taco1* was induced using double-stranded dicer substrate (dsiRNA, IDT). Efficacy of each siRNA was determined experimentally during and after differentiation. For *Slc25a3*, siRNAs *Slc25a3*.*13*.*2* (targeting 3’ UTR) and *Slc25a3*.*13*.*1* (targeting exon 4) were used at a concentration of 50 µM per well in differentiating myoblasts and 150 µM in differentiated myotubes. Specificity of dsiRNA was validated by loss of SLC25A3 protein and rescue with expression of wild type SLC25A3 protein. For *Cox17*, dsiRNA *Cox17*.*13*.*3* (targeting exons 1,2) was used at a concentration of 50 µM per well in differentiated myoblasts and differentiated myotubes. Specificity of dsiRNA was validated by loss of COX17 protein. For *Taco1*, dsiRNA *Taco1*.*13*.*2* (targeting exon 1) was used at a concentration of 50 µM per well in differentiating myoblasts, though no concentration tested induced effective TACO1 protein loss in myotubes. Specificity of dsiRNA was validated by loss of TACO1 protein in differentiated myoblasts. The dsiRNAs or corresponding negative control non-targeting dsiRNAs (IDT 51-01-14-04) were introduced by transfection using Lipofectamine 3000 (Invitrogen, L3000008) according to the manufacturer’s instructions as previously described (37). For experiments to induce knockdown in differentiating myoblasts, C2C12 or primary cells plated for normal proliferation, incubated with transfection mixes containing gene targeting or negative control dsiRNA in growth medium without antibiotics overnight. After 16– 18 h, transfection medium was replaced with differentiation medium (C2C12), or cells were plated for differentiation on coated plates (primary) and appropriate concentrations of BCS or CuSO_4_ were added. For experiments with knockdown in myotubes, cells were differentiated until myotubes were visible and then myotubes were incubated with transfection mix containing gene targeting or negative control dsiRNA in differentiation medium for 16-18 hours before being returned to differentiation medium. In all cases, cells were harvested 48 hours after transfection. For experiments with co-expression of SLC25A3 isoforms or variants, plasmids encoding *Slc25a3* were co-transfected with dsiRNAs.

### Immunoblotting

To harvest for immunoblotting, cells were washed twice in phosphate buffered saline (PBS, Fisher Bioreagents BP665-1) and incubated in radioimmunoprecipitation buffer [RIPA: 10 mM piperazine-*N*-*N*-bis (2-ethanesulfonic acid)] supplemented with protease inhibitor (Pierce, PIA32953) and phosphatase inhibitor (Sigma Aldrich, p2850). Samples were sonicated on ice two times, 10 seconds each at 40% output on an ultrasonic membrane disruptor (Fisher Model 100). After sonication, samples were incubated on ice for 30 min and centrifuged at 21,000× *g* for 30 min. The supernatant was transferred to new tubes and total protein concentration was determined using Bradford assay (Bio-Rad 5000205). To prepare for SDS-PAGE, protein was diluted in Laemmli sample buffer (Bio-Rad 161-0747). For soluble proteins, samples were boiled for 5 minutes, and samples were not boiled for membrane bound proteins. Proteins were separated on TGX stain-free 4–20% polyacrylamide gels (Bio-Rad 4568094) and then transferred to nitrocellulose membrane (Cytiva, 45-004-001) using a Trans-Blot Turbo semi-dry transfer system (Bio-Rad) or in a wet transfer apparatus for 1 hour kept cold with agitation. Unless otherwise noted, gels were loaded with 20 µg of lysate. After transfer, blots were blocked in Every Blot blocking reagent (Bio-Rad 12010020). Blots were probed with primary antibodies for proteins of interest (Supplemental Table 1) overnight at 4°C with gentle agitation and then probed with horseradish peroxidase (HRP)-conjugated secondary antibodies for 1 hour at room temperature. Target proteins were visualized using Pico Plus (ThermoFisher PI34580) or Femto Plus (ThermoFisher PI34095) ECL substrate on a Bio-Rad Chemi-doc imager. In all cases, Ponceau staining to visualize total protein was used as a loading control.

### Immunofluorescence staining and quantification

Immunofluorescence staining for eMyHC was used to visualized myotubes as previously described (37). Briefly, cells were washed in PBS and fixed with 3.7% formaldehyde or 4% paraformaldehyde for 10 minutes at room temperature followed by three additional washes in PBS. Cells were incubated in blocking buffer, 3% BSA with 0.3% Triton X-100 (Fisher, BP151 100), for 1 hour at room temperature and incubated with an antibody to eMyHC overnight at 4°C. The following morning, cells were washed three times with 0.5x blocking buffer and incubated with FITC-conjugated anti-mouse antibody in 0.5x blocking buffer for 1 hour at room temperature protected from light. Cells were washed three times with 0.5x blocking buffer and stained with 4′-6-diamidino-2-phenylindole (DAPI, Fisher D3571) diluted 1:1000 in PBS for 5 minutes. Cells were washed twice in PBS and incubated in fresh PBS for imaging. Cells were imaged using an Olympus IX83 inverted fluorescence microscope using a U Plan fluorite 10X phase objective lens (NA 0.3 WD 10 mm). Images were captured using a DP74 Color CMOS Camera (cooled 20.8 MP pixel-shift, 60 FPS) using CellSens Dimension V2 software. Images were subjected to 2D-deconvolution and exported as red-green-blue (RGB) tif files. For each experimental condition, three images were captured at random areas in each well. For representative images shown in figures, linear level adjustments were performed in Adobe Photoshop evenly across all images to aid in visualization. All adjustments were performed in separate designated files after any quantitative measurements were collected.

All quantitative analysis was performed by an individual blinded to the identity of the sample shown in the image on three technical replicates per each biological replicate. Fusion index was quantified as described wherein nuclei in eMyHC positive multinucleated (≥2 nuclei/tube) myotubes were counted and normalized to total DAPI-stained nuclei (37). For total eMyHC fluorescence and live/dead staining, total green fluorescence was measured and normalized to total blue fluorescence (DAPI/Hoechst) using FIJI. First, green and blue channels were separated, and each image was subjected to threshold adjustment and de-speckling as needed to minimize background signal. Total green and blue fluorescence was then measured using the analyze integrated density function for the entire image.

### Cell viability assays

Cell viability assays were performed using RealTime-Glo MT Cell Viability Assay (Promega G9711) according to the manufacturer’s instructions for endpoint assay. Briefly, myotubes were differentiated in white clear-bottom 96 well plates until controls contained fully differentiated myotubes. On the day of the assay, enzyme and substrate solutions were mixed in differentiation medium, warmed to 37°C, and used to replace differentiation medium on cells. After 30 minutes incubation at 37°C, bioluminescence was imaged on a BioTek Gen5 plate reader. After cells were assayed, they were harvested in RIPA buffer, sonicated as described above, and subjected to Bradford analysis for total protein concentration. Cell viability was then calculated as luminescence/ protein concentration as determined by Bradford assay. For cell permeability-based viability assay, the ReadyProbes cell viability imaging kit (ThermoFisher R37609) was used according to the manufacturer’s instructions. Cells were provided with 2 drops each of NucBlue Live and NucGreen Dead stain, incubated for ~20 minutes, and imaged as described above. Total live and dead cells were quantified by measuring total blue and total green fluorescence as described above.

### Molecular cloning

The DNA sequences for *Slc25a3* isoforms A, B and variants were synthesized and purchased from Twist Bioscience. They were then subcloned into a pcDNA3.1 vector using double digest using restriction endonucleases Not1 and Xho1 enzymes. All constructs were confirmed by diagnostic digest, Sanger Sequencing (Azenta Bioscience) and whole plasmid sequencing (Plasmidsaurus).

### Alpha fold modeling of SLC25A3 structure

Sequences for murine SLC25A3-A and -B were translated from ENSMUST00000170810.8 for *Slc25a3-A* and ENSMUST00000076694.13 for *Slc25a3-B*. Sequences were modeled in AlphaFold 3 (75).

### RNA isolation, reverse transcription, and quantitative PCR

Steady-state levels of transcripts were determined using quantitative PCR on reverse-transcribed cDNAs (RT-qPCR) as previously described (37). Total RNA was isolated with TRIzol (Invitrogen, 15596018) according to the manufacturer’s instructions and cDNA was synthesized from 1 µg of RNA using the Maxima First Strand cDNA Synthesis Kit with dsDnase treatment for 30min (ThermoFisher, K167). Quantitative PCR was performed using SYBR Select Master Mix (Applied Biosystems, 4472908) on a QuantStudio 3 Real Time PCR system (Applied Biosystems). All primers were designed using NCBI Primer-Blast and sequences can be found in Supplemental Table 2. Results were quantified using the comparative Ct method with *Rplp0* used as a normalizer (76).

### Electron flow assays

Seahorse XF Pro Analyzer (Agilent) was used to perform electron flow assays using mitochondrial respirometry from frozen cell stocks as previously described (41). Cells were plated and harvested at myoblast stage or differentiated to myocyte and myotube stages prior to harvest. At harvest, cells were rinsed in PBS and then scraped and pelleted in PBS before flash freezing in liquid nitrogen. Frozen cell pellets were thawed on ice and suspended in MAS buffer (220 mM Mannitol, 70 mM Sucrose, 10 mM KH_2_PO_4_, 5 mM MgCl_2_, 1 mM EGTA, 2 mM HEPES pH7.2) and homogenized in bead mill at 0°C with a speed of 6 m/sec for 30 sec. Homogenate is centrifuged at 900 x g for 10 min at 4°C and clear supernatant is used for protein estimation with BCA (Thermo Scientific 23225) method. Frozen respirometry assay is performed with 10ug protein per reaction in assay buffer containing 10 ug/mL cytochrome C in MAS buffer. With sequential injections of either 1 mM NADH (Complex I substrate) or 5 mM Succinate with 2 µM Rotenone (Complex II substrate and Complex I inhibitor respectively) in port A, 4 µM Antimycin (Complex III inhibitor) in port B, 0.5 mM TMPD with 1 mM Ascorbate (Complex IV substrate) in port C and 50 mM sodium azide (Complex IV inhibitor) in port D, oxygen consumption (OCR) is measured and normalized with total protein content taken per reaction.

### ATP rate assays

ATP rate assays were performed as described (77). Myoblasts were plated in an XF cell culture microplate at time 0 and changed to differentiation medium (myotube experiments). For myoblast experiments, cells were plated later once myotube formation was nearly complete in myotube experiment wells. Once both cell populations were confluent or differentiated into myotubes ATP rate assays were performed on both populations. Seahorse XF DMEM medium was prepared with 10mM XF glucose, 1mM XF pyruvate, and 2mM XF glutamine. Cells were washed in Seahorse XF medium and incubated for one hour. XF ATP Rate Assay (Agilent #103592–100) was performed according to the manufacturer’s instructions on a Seahorse XFe96 Analyzer (Agilent #S7800B). Data were exported and analyzed using the Seahorse Wave Desktop Software (Agilent).

### Statistical analyses

All statistical analysis was performed using GraphPad Prism version 11.0.1 for MacOS. Quantification in figures show mean ± standard error of the mean. As noted, all quantifications comparing one variable in two groups used paired two-tailed *t* testing unless otherwise noted, quantifications comparing three or more groups used paired one-way ANOVA with Dunnett’s post hoc correction for multiple comparisons, and quantifications comparing more than one related variable in two or more groups, two-way ANOVA with Sidak’s post hoc correction for multiple comparisons. In all cases, p < 0.05 was considered statistically significant. For all quantifications, number of biological replicates, statistical tests, and ranges of p values are included in figure legends.

## Data availability

All data are contained within the manuscript.

## Supporting information

This article contains supporting information.

## Acknowledgements

The authors would like to thank Dr. Paul Cobine (Auburn University) and Dr. Scot Leary (University of Saskatchewan) for generously sharing the custom antibody to SLC25A3 (45).

## Figures and legends

**Supplemental Figure 1:**
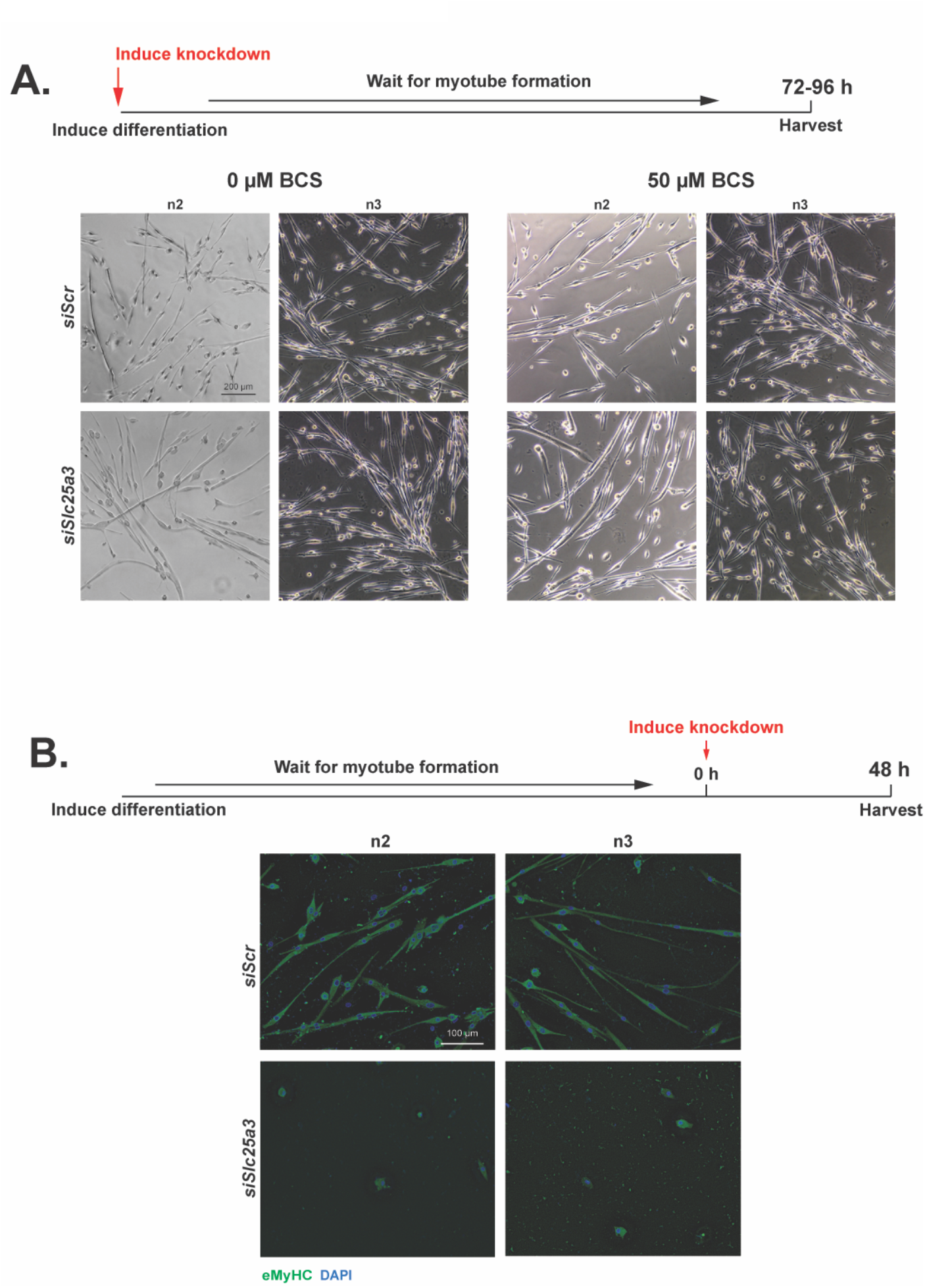
Additional replicates of SLC25A3 deficient primary myotubes. *A)* Schematic of experiment wherein siRNA was used to knock down *Slc25a3* at the same time that differentiation was induced. *B)* Representative phase contrast images for additional replicates of primary myotubes showing no overt defect in myotube formation caused by *siSlc25a3* with or without addition of 50 µM BCS. Shown are images from n=2 additional biological replicates. Bar=200 µm. *C)* Schematic of experiment wherein myoblasts were fully differentiated to myotubes and then siRNA used to knock down *Slc25a3* with or without BCS for 48 hours. *D)* Representative immunofluorescence images for additional replicates of primary myotubes showing cell death in myotubes caused by *siSlc25a3*. Myotubes were stained with an antibody eMyHC and DAPI to visualize nuclei. Shown are images from n=2 additional biological replicates. Bar=100 µm.

**Supplemental Figure 2:**
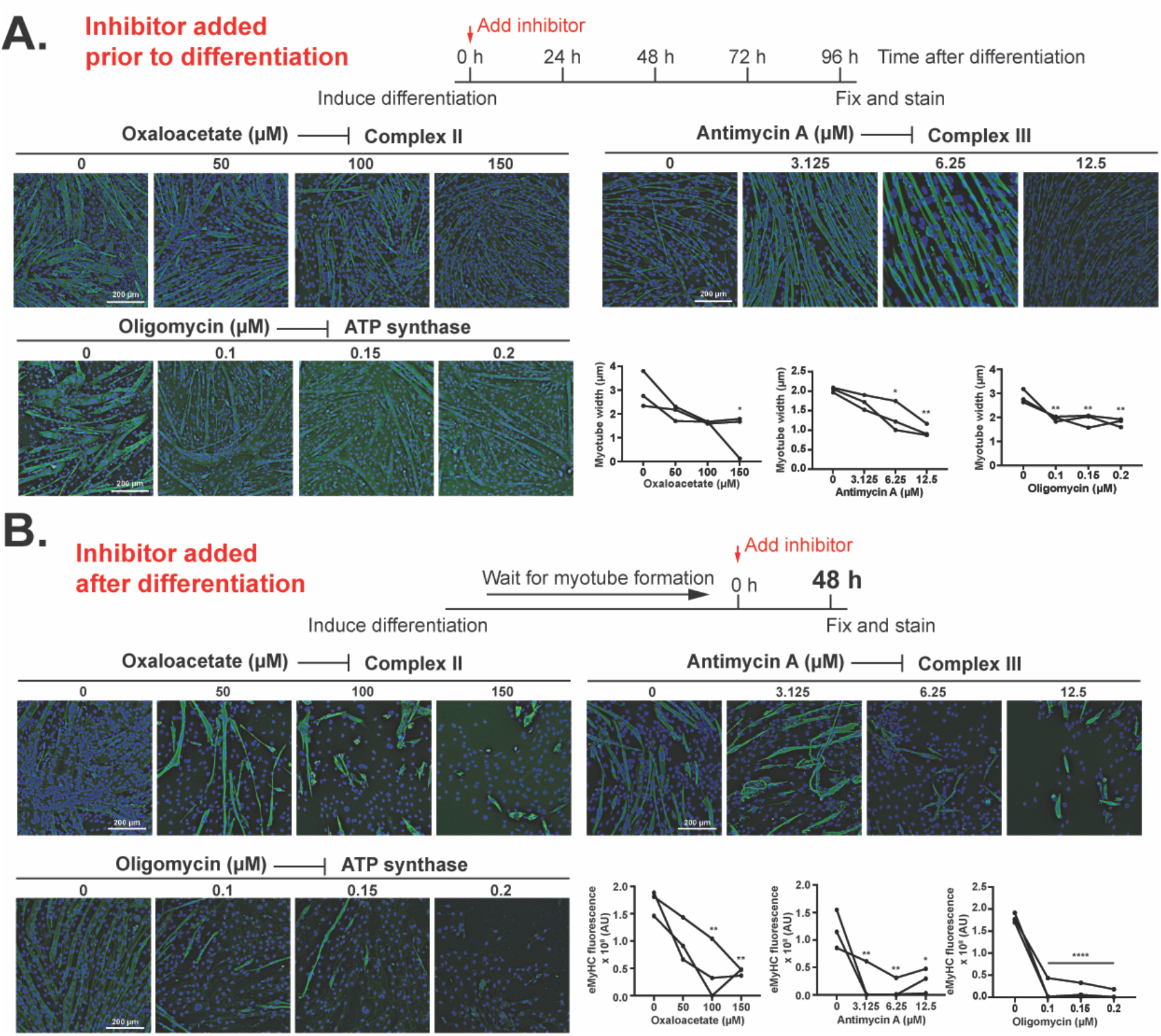
Chemical inhibition of electron transport chain complexes II, III or ATP synthase impairs myotube survival but not formation. A) Representative images and quantification of myotube width showing that addition of inhibitors to electron transport chain complexes II (oxaloacetate), III (antimycin a), and ATP synthase (oligomycin) at the same as inducing myoblast differentiation does not inhibit myotube formation but results formation of smaller myotubes. *B)* Representative images and quantification of eMyHC fluorescence for fully differentiated myotubes treated with inhibitors to complexes II, III, and ATP synthase as in *A*. For both *A* and *B*, immunofluorescence images show myotubes stained with an antibody eMyHC and DAPI to visualize nuclei. For quantification, shown are individual data points for n=3 biological replicates. Statistical significance was determined using one-way ANOVA with Dunnett’s post-hoc test. *p<0.05, **p<0.01, ****p<0.0001.

**Supplemental Figure 3:**
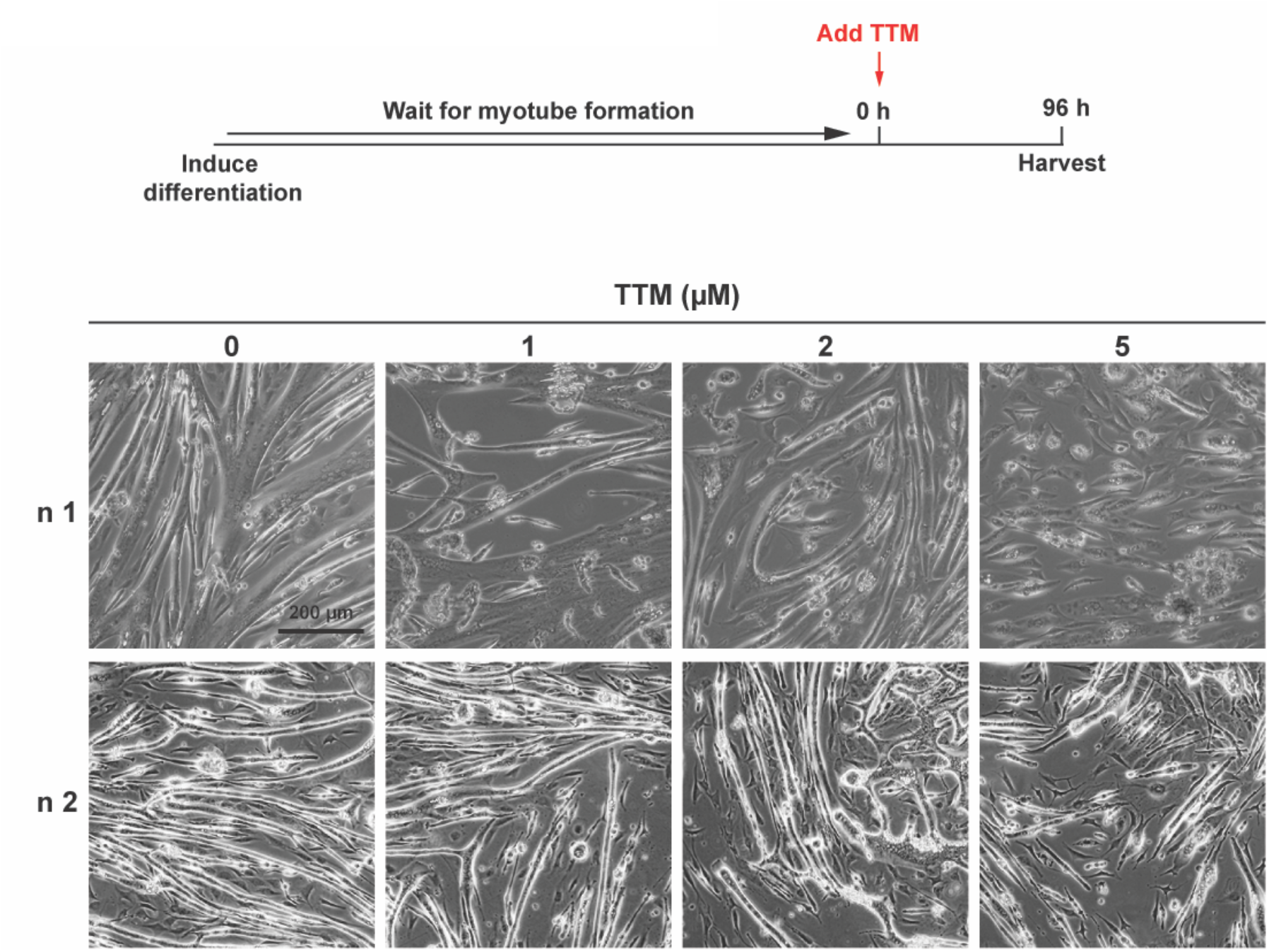
Copper chelation in fully differentiated myotubes leads to detachment and death. Schematic of experiment showing that tetrathiomolybdate (TTM) was added to fully differentiated myotubes for 96 hours. Representative phase contrast images showing detachment and death of myotubes treated with TTM in a dose-dependent manner.

**Supplemental Table 1:**
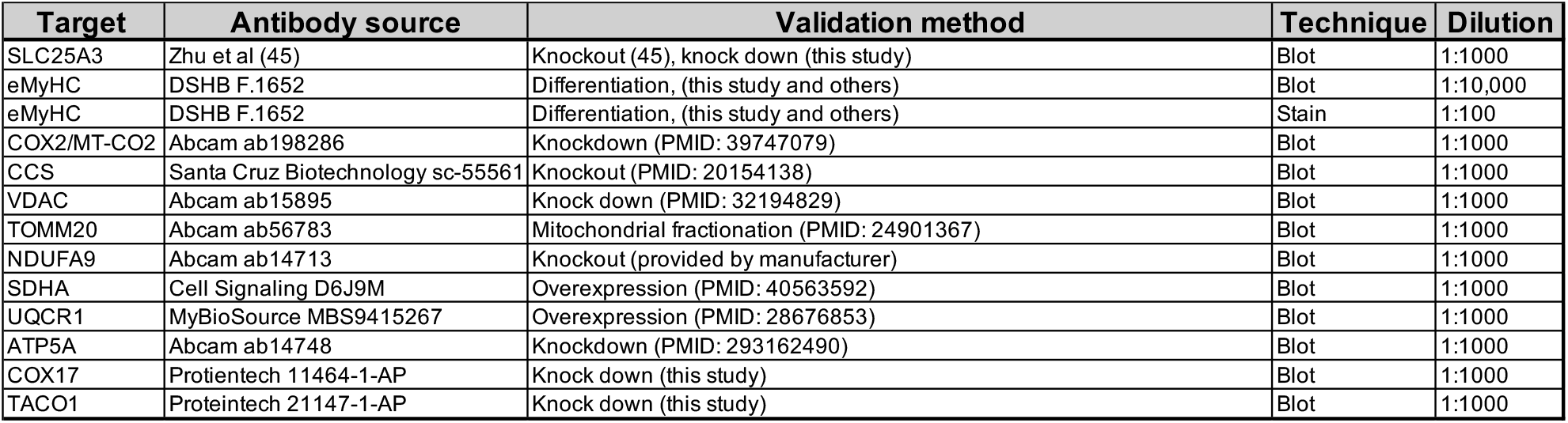
Detailed antibody information for this study.

**Supplemental Table 2:**
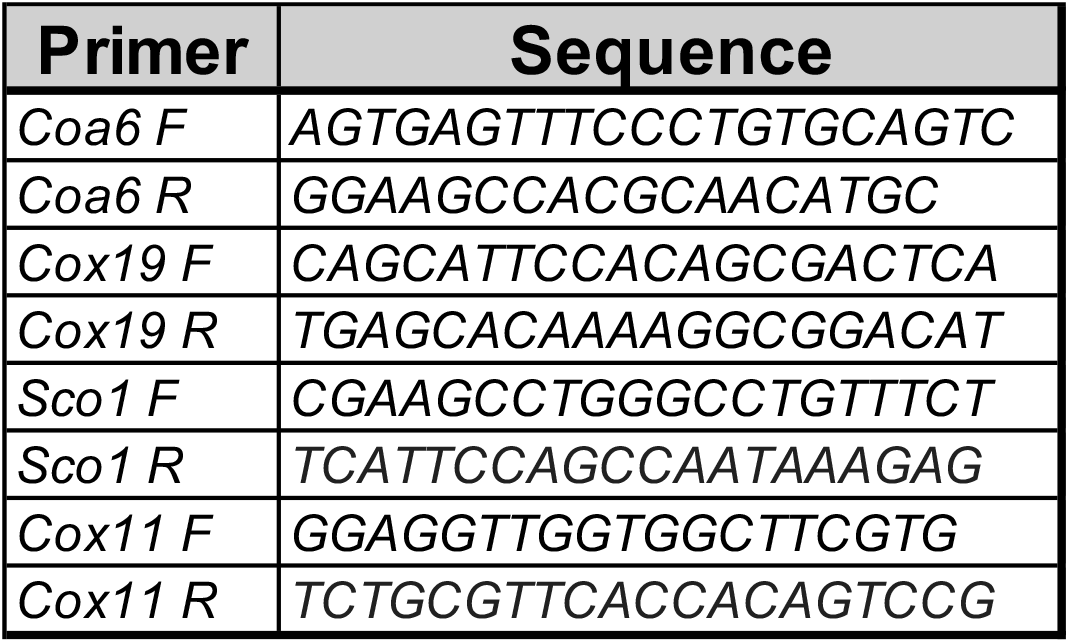
Primers for qRT-PCR used in this study.

