## Supplemental data for "Copper transport to mitochondria by SLC25A3 contributes to skeletal myoblast differentiation and is required for survival of differentiated myotubes"

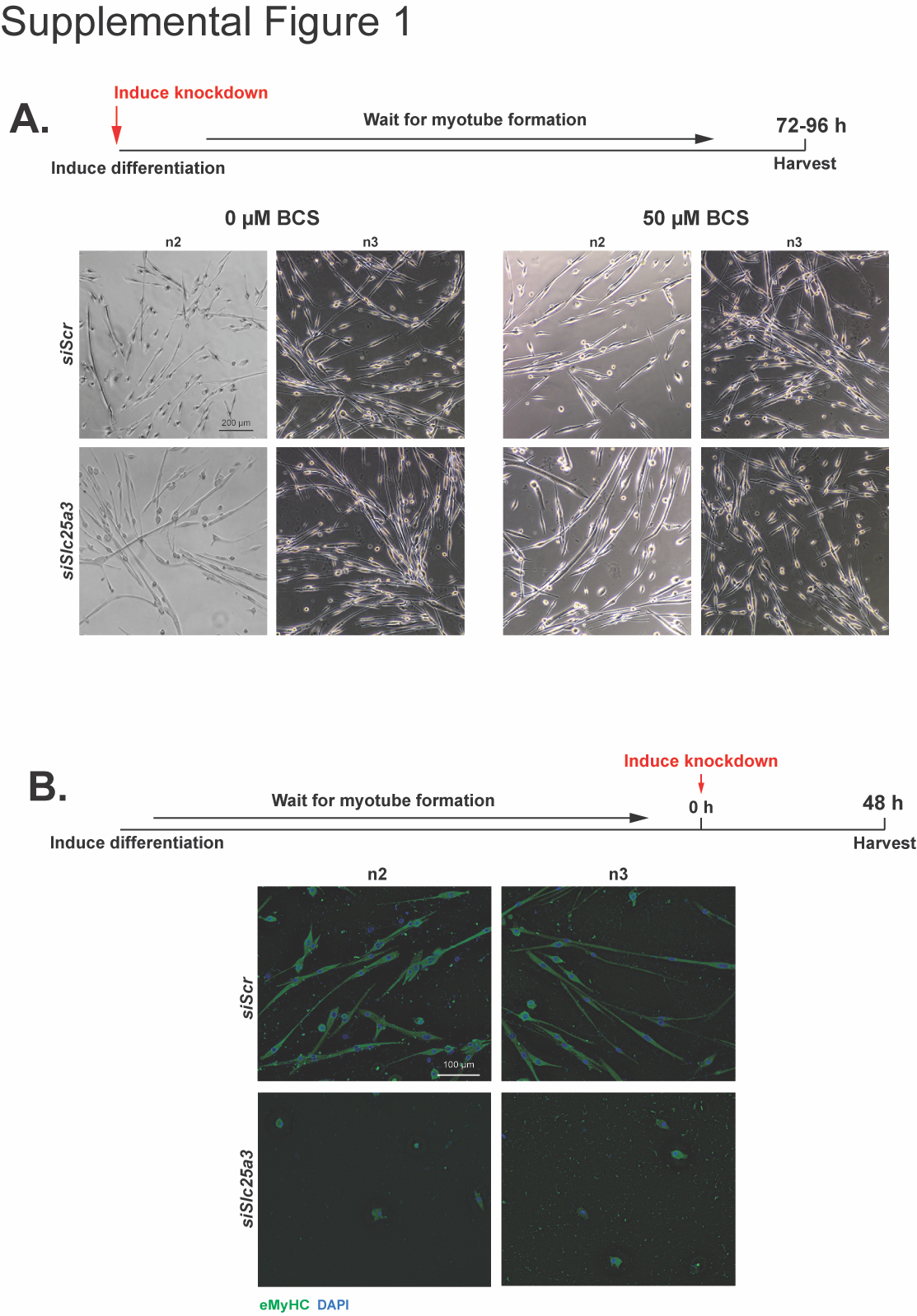

**Supplemental Figure 1:** **Additional replicates of SLC25A3 deficient primary myotubes.** *A)* Schematic of experiment wherein siRNA was used to knock down *Slc25a3* at the same time that differentiation was induced. *B)* Representative phase contrast images for additional replicates of primary myotubes showing no overt defect in myotube formation caused by *siSlc25a3* with or without addition of 50 µM BCS. Shown are images from n=2 additional biological replicates. Bar=200 µm. *C)* Schematic of experiment wherein myoblasts were fully differentiated to myotubes and then siRNA used to knock down *Slc25a3* with or without BCS for 48 hours. *D)* Representative immunofluorescence images for additional replicates of primary myotubes showing cell death in myotubes caused by *siSlc25a3*. Myotubes were stained with an antibody eMyHC and DAPI to visualize nuclei. Shown are images from n=2 additional biological replicates. Bar=100 µm.

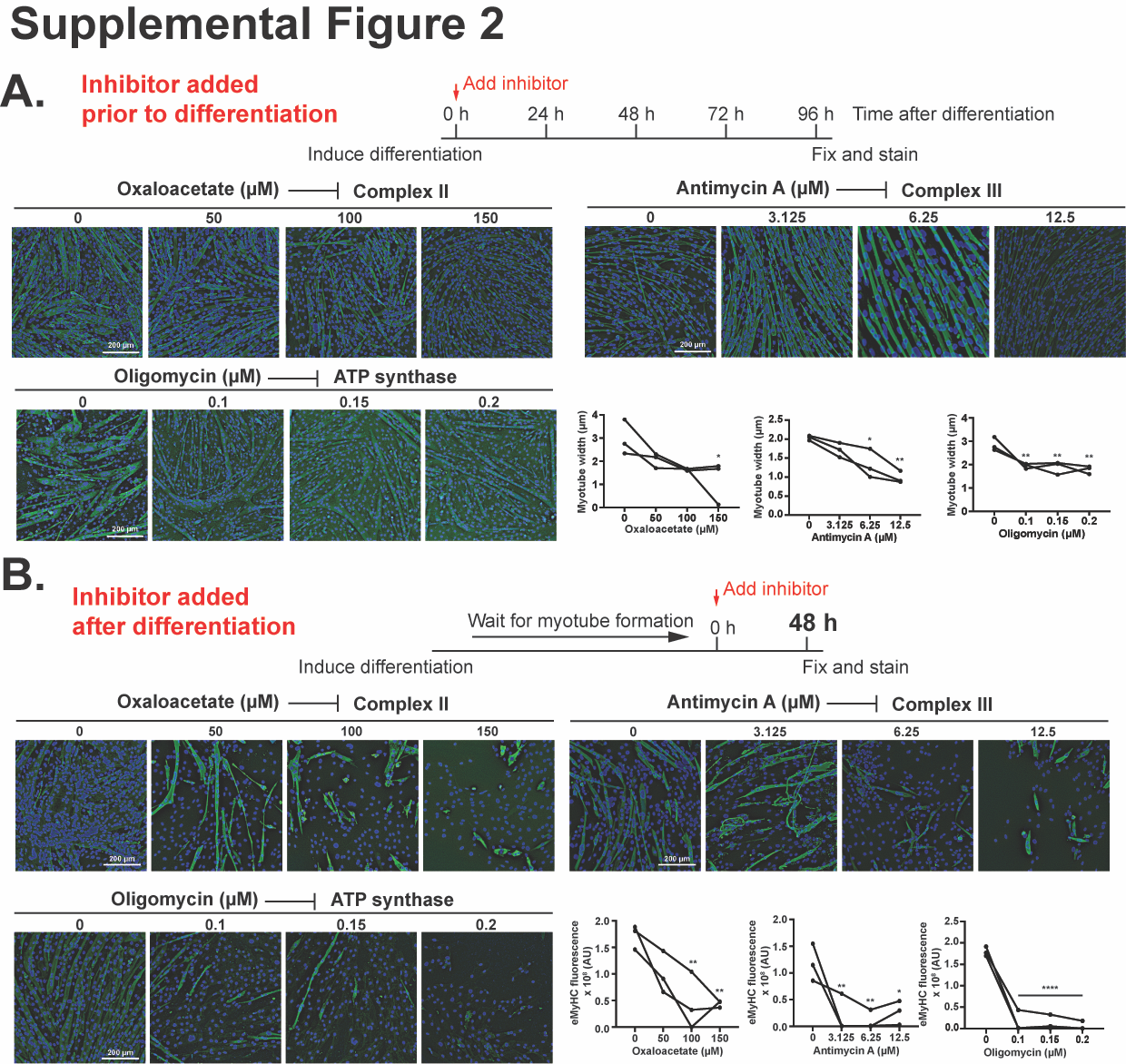

**Supplemental Figure 2: Chemical inhibition of electron transport chain complexes II, III or ATP synthase impairs myotube survival but not formation.** A) Representative images and quantification of myotube width showing that addition of inhibitors to electron transport chain complexes II (oxaloacetate), III (antimycin a), and ATP synthase (oligomycin) at the same as inducing myoblast differentiation does not inhibit myotube formation but results formation of smaller myotubes. *B)* Representative images and quantification of eMyHC fluorescence for fully differentiated myotubes treated with inhibitors to complexes II, III, and ATP synthase as in *A*. For both *A* and *B*, immunofluorescence images show myotubes stained with an antibody eMyHC and DAPI to visualize nuclei. For quantification, shown are individual data points for n=3 biological replicates. Statistical significance was determined using one-way ANOVA with Dunnett’s post-hoc test. *p<0.05, **p<0.01, ****p<0.0001.

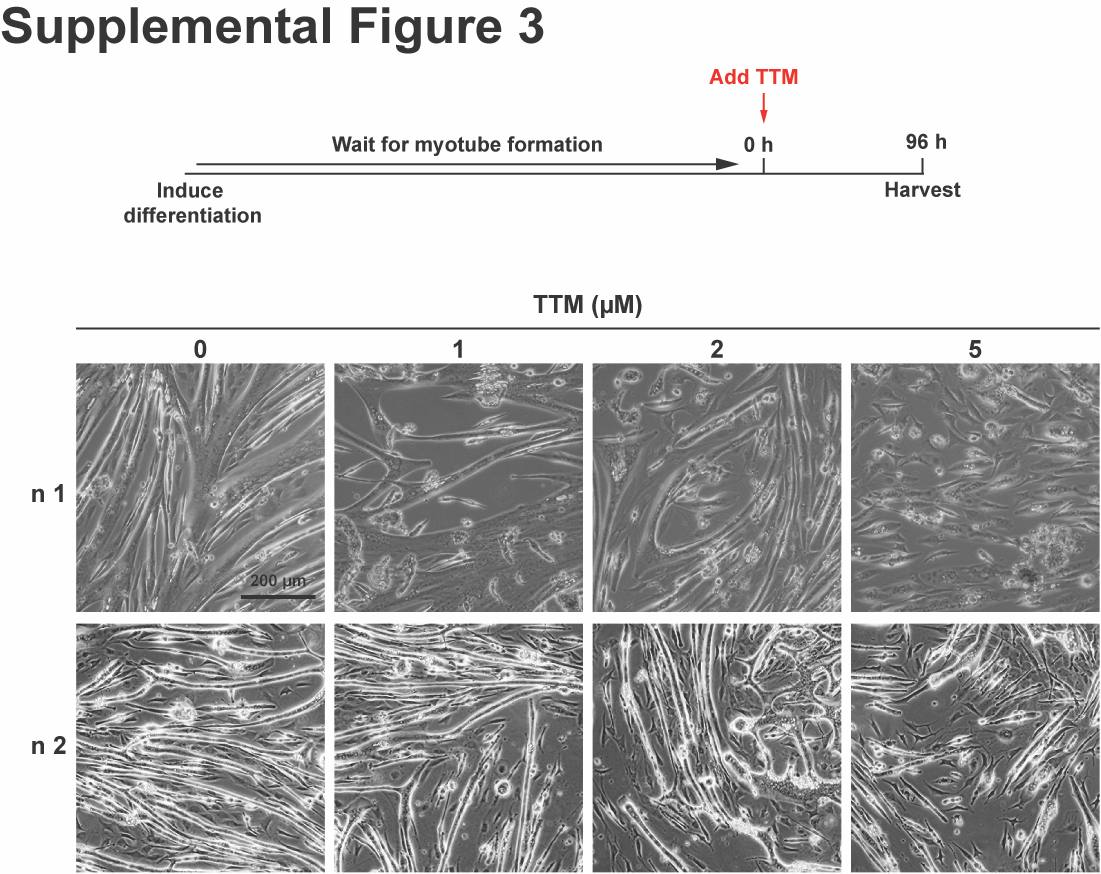

**Supplemental Figure 3: Copper chelation in fully differentiated myotubes leads to detachment and death.** Schematic of experiment showing that tetrathiomolybdate (TTM) was added to fully differentiated myotubes for 96 hours. Representative phase contrast images showing detachment and death of myotubes treated with TTM in a dose-dependent manner.

**Supplemental Table 1: Detailed antibody information for this study**

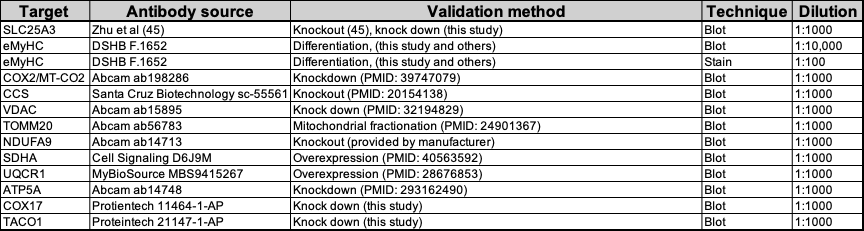

**Supplemental Table 2: Primers for qRT-PCR used in this study**

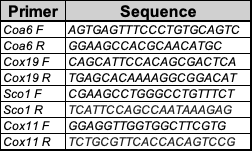
